# Lectin Microarray-based Glycomics and Machine Learning Identify Shared Osteoarthritis Biomarkers in Humans, Dogs, and Horses

**DOI:** 10.1101/2025.10.16.682971

**Authors:** Angelo G Peralta, Parisa Raeisimakiani, Kei Hayashi, Lara K Mahal, Heidi L Reesink

## Abstract

Post-traumatic osteoarthritis (PTOA) is a common sequela to joint injury in both humans and companion animal species such as horses and dogs. Despite the increasing prevalence of osteoarthritis (OA) in humans, investigation of glycosylation changes associated with OA remains in its infancy. Recent advances, such as lectin microarray analysis, now enable detailed glycan profiling in complex biofluids such as synovial fluid. Using lectin microarray technology, this study characterized glycosylation patterns in synovial fluid samples from healthy and OA-affected joints in horses, dogs, and humans. Comparative glycan-binding profiles within and between species revealed conserved and distinct glycomic signatures associated with OA. Machine learning models, including classification algorithms, effectively distinguished OA from healthy joints, identifying key lectins and glycan epitopes crucial to these predictions. The identified lectin markers reflect specific glycosylation pathways and potential inflammatory mechanisms, demonstrating their value in differentiating between healthy and OA phenotypes. Our findings underscore the promise of integrated glycomic profiling and machine learning to enhance our understanding of glycan involvement in the pathogenesis of OA and to facilitate the development of diagnostic and therapeutic strategies applicable to both veterinary and human medicine.

**In Brief:** Osteoarthritis affects humans and companion animals; however, its molecular features remain unclear. Using lectin microarrays and machine learning, we identified conserved and species-specific glycan signatures in synovial fluid that differentiate between control and osteoarthritic joints. This One Health approach highlights shared molecular mechanisms of joint degeneration and establishes data-driven glycomic profiling as a framework for understanding osteoarthritis across species.

**Graphical Abstract:** 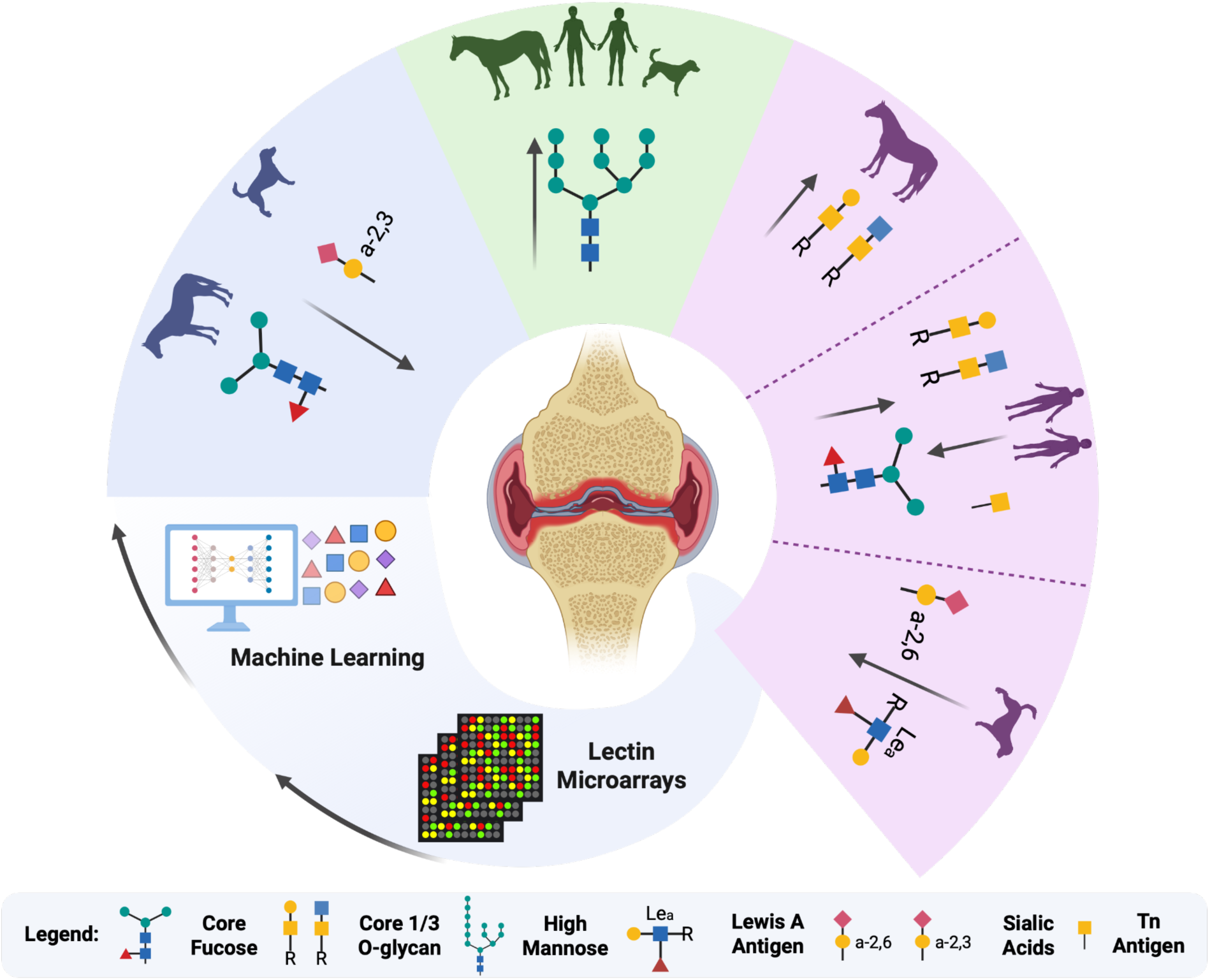

## Introduction

Post-traumatic osteoarthritis (PTOA) occurs secondary to joint injury, such as an intra-articular fracture, ligament tear, or cartilage impact injury and is characterized by pain, stiffness, and impaired joint function and mobility. Osteoarthritis (OA) results in significant morbidity in both humans and domestic veterinary patients. The prevalence of OA is rising sharply with global aging, with PTOA accounting for approximately 12.4% of all symptomatic OA cases (1, 2). Twenty percent of dogs older than one year and up to 80% of geriatric dogs are diagnosed with osteoarthritis, and lameness due to joint disease or osteoarthritis is the leading reason that horses present to equine veterinarians.(3–5). While the underlying mechanisms of OA are under active investigation, the lack of early diagnostic methods hamper early intervention, leaving current approaches focused on palliating symptoms through medication or surgery rather than slowing disease progression (6–8). Although radiography is widely used for OA diagnosis, it lacks the sensitivity to detect early pathological changes and correlates poorly with joint function(6, 9). Given these limitations and the restricted use of more sensitive imaging modalities such as MRI or CT contrast arthrography, there is a need for minimally invasive biomarkers to enable earlier diagnosis and better disease monitoring.

Synovial fluid (SF) is a joint-derived biofluid that provides a highly localized molecular snapshot of osteoarthritis (OA) pathophysiology due to its direct contact with articular cartilage, synovium, ligaments, and other affected tissues (10, 11). This spatial proximity enables SF to reflect early and dynamic biochemical changes more accurately than peripheral biofluids such as serum, plasma, or urine (12). Metabolomic profiling has demonstrated significant alterations in osteoarthritic SF, supporting its utility in biomarker discovery. For example, metabolomic analysis detected alterations in SF between healthy and PTOA equine carpal joints, including changes related to inflammation, oxidative stress and collagen synthesis and degradation pathways, underscoring its potential for early OA detection (13, 14). While proteomic and metabolomic approaches have identified candidate biomarkers and glycoproteomic analyses have begun to reveal novel diagnostic targets, a systematic interrogation of SF glycan structures across species has not been performed to date (11, 13, 15, 16).

Glycans, which can be present as free glycans or attached to proteins and lipids, play essential roles in regulating biological processes in both health and disease (17).

Structural alterations in glycans—referred to as dysregulated glycosylation—have been associated with a wide range of conditions, including immune disorders, infectious diseases, and cancer (18–20) Characterizing the glycome, or the complete set of glycans expressed in a biological system, is therefore critical to understanding the molecular basis of health and disease (21).

Recent advances in high-throughput glycomic technologies have begun to overcome these limitations, enabling the rapid and comprehensive profiling of glycan structures (22, 23). The increasing availability of large-scale glycan datasets (24), particularly from analytical-based (22) and array-based platforms (25, 26), provides new opportunities for computational analysis. In this context, machine learning methods offer powerful tools for extracting hidden patterns and functional insights from complex glycomic data (27, 28), helping to advance biomarker discovery (29) and deepen our understanding of glycan-mediated mechanisms in disease (30).

In this study, SF samples were collected from equine, canine, and human joints and categorized as control or osteoarthritic. Glycosylation profiles were analyzed using lectin microarray technology to assess differences in SF glycopatterns both within and across species. By integrating statistical analysis with machine learning, we evaluated the potential of SF glycopatterns as cross-species diagnostic indicators of OA. This study provides novel insights into conserved glycosylation changes across species and supports the feasibility of using SF glycomics for early diagnosis and patient stratification in OA. Our work advances glycomic analysis of synovial fluid in a way that brings glycan-based biomarkers closer to practical application for OA diagnosis and treatment. Finally, the multi-species, multi-method approach presented here highlights the potential for glycan biomarkers to inform future diagnostic and therapeutic strategies for OA in both humans and veterinary patients.

## Results

### Sample demographics

The human cohort consisted of 52 participants: 25 with full anterior cruciate ligament (ACL) rupture and 11 with advanced osteoarthritis (OA) of the knee necessitating total joint arthroplasty (TKR). The remaining 16 contralateral knees (10 from the ACL group and 6 from the TKR group) acted as controls. Importantly, controls were not recruited as separate healthy individuals; rather, they represent the contralateral (non-injured/non-arthritic) knees from the same participants who presented for ACL or TKR procedures. PTOA cases were younger on average (median: 35.5 years, IQR: 37) compared with contralateral controls (median: 49.5 years, IQR: 36.5). Males and females represented 50% and 50% of the ACL injury group and 66.7% and 33.3% of the TKR group, respectively (see Table S1 for complete demographic details).

The equine cohort included 118 horses, with 77 affected by PTOA and 41 classified as healthy controls. Horses in the PTOA group were generally younger (median 3 years, IQR: 3 years) than controls (median 5 years, IQR: 2.4 years). In the control group, the sex distribution was 51% females, 41% castrated males, and 7% intact males while the PTOA group consisted of 46% females, 41% castrated males, and 13% intact males.

While the majority of PTOA cases involved the carpus (middle carpal joint [MCJ] or antebrachiocarpal joint [ACJ]), SF samples were included from other PTOA joints, including the fetlock (metacarpo-/metatarsophalangeal joint [MCPJ/MTPJ]), tibiotarsal joint, and knee (stifle joint) (see Table S2 for full demographic details).

The canine cohort consisted of 60 subjects, including 51 dogs with PTOA of the stifle associated with cranial cruciate ligament rupture and 9 healthy controls. Median age was greater in the PTOA group (6 years, IQR: 4.4 years) than in controls (4 years, IQR: 1.5). All control dogs were healthy males, whereas the PTOA group included a mix of neutered males (50%), spayed females (48%), and one intact female (2%) (see Table S3 for complete demographic details).

### Diagnostic potential of synovial fluid glycomics

While glycosylation changes have been examined in certain joint diseases, such as rheumatoid arthritis (31, 32) and spondyloarthropathies (33, 34), their role in PTOA remains poorly understood (35–41). To investigate whether glycan profiles in synovial fluid could distinguish healthy from OA-affected joints across species, we employed lectin microarray profiling. Lectin microarrays enable the identification of disease-associated glycan signatures in complex biofluids (38, 42–44). Thus, we performed lectin microarray analysis using a dual-color labeling strategy, where each sample was labeled with a fluorophore and hybridized against a pooled reference mixture labeled with an orthogonal fluorophore (26, 45). Lectin microarrays utilize carbohydrate-binding proteins with defined glycan specificities to detect glycan epitopes, enabling a systems-level view of the glycome (22, 46, 47). In this study, 62 probes were used (detailed in the supporting information, Data S1) and epitopes were annotated using literature (48–50). Differential glycomic epitope P-values < 0.05 were extracted, displayed with heat maps **(Figure S1A-C)**, and evaluated using volcano plots **(Figure 1-3)**.

**Figure 1:**
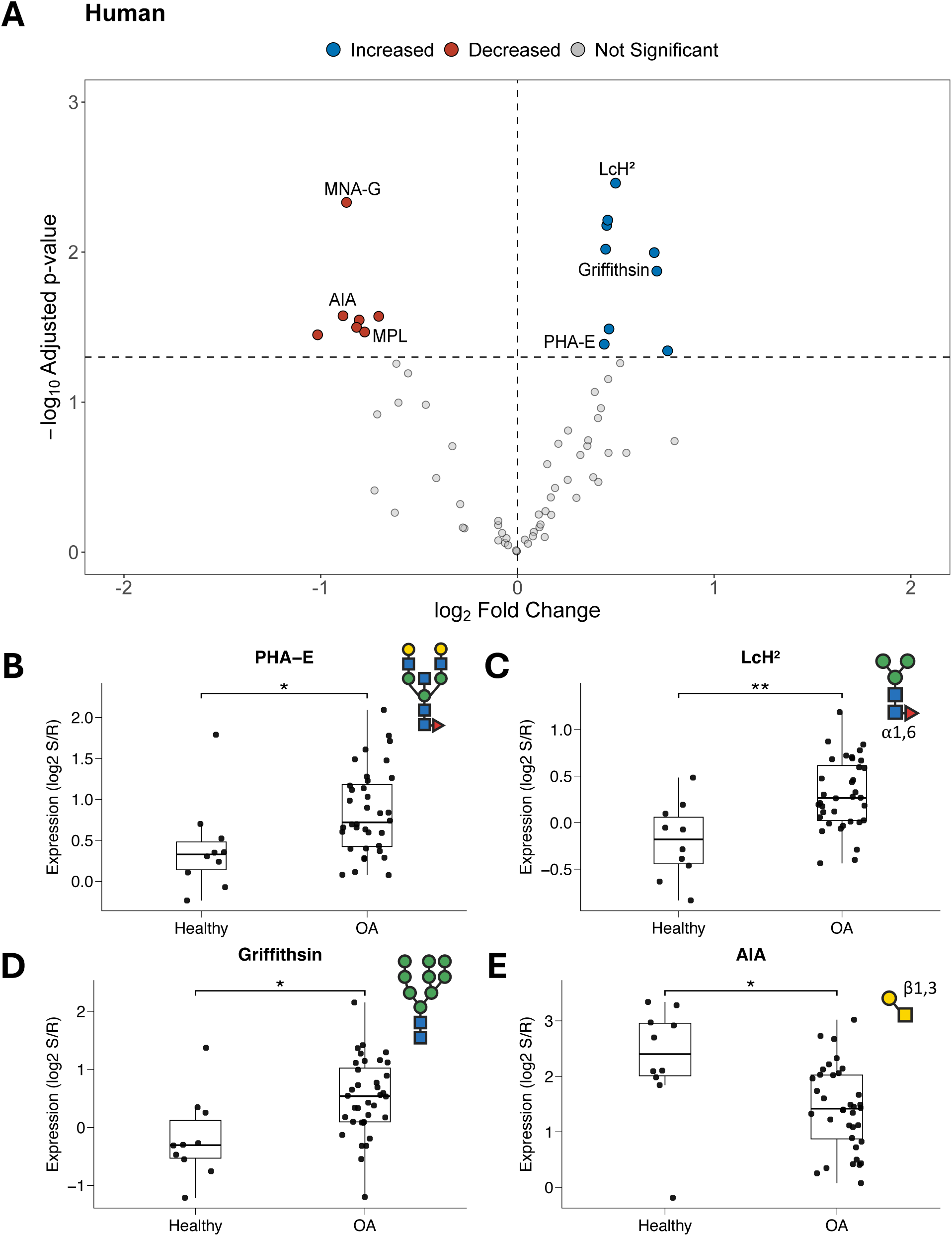
Glycomic alterations in human synovial fluid associated with osteoarthritis (OA). (A) Volcano plot comparing lectin microarray signal intensities between OA and control subjects. Blue and red dots represent lectins with significantly lower or higher signal intensities, respectively (FDR-adjusted p < 0.05). (B–E) Selected lectins recognizing branched N-glycans, core fucosylated glycans, high-mannose glycans, and core 1/3 O-glycans show altered binding in OA synovial fluid. p < 0.05, *p < 0.01, \*\**p* < 0.001. Lectin labels are shown with abbreviated names; duplicated probes are distinguished by numeric superscripts (e.g., LcH¹, LcH²; anti-LeA¹, anti-LeA²). See Table S4 for full description.

### Species-specific glycan alterations in OA

Our lectin microarray data revealed glycan alterations at three levels: changes shared across all three species, changes common to horses and dogs, and changes unique to individual species. In canine OA joints, significant decreases in both α2,6-and α2,3-sialylation were detected, as indicated by reduced binding of SNA, PSL1a, SLBR-N, and dicCBM40 **(Figure 3A)**. Decreased lectin binding signals for Lewis A antigens and core fucosylation (LcH, PSA) were also specific to the canine OA samples **(Figure 3A**, **3E)**. In comparison, type 3/4 H-antigens (TJA-II, SNA-II) were elevated in canine OA joints compared to controls **(Figure 3A)**. Equine OA joints exhibited distinct increases in core 1/3 *O*-glycans (AIA, MPL) **(Figure 2A, 2B)** which were not observed in human or canine samples **(Figure 1A, 3A)**. These glycomic changes suggest a mucin-related response in the equine joint, while alterations in sialylation were observed in the canine cohort. In human OA joints, several unique lectin profiles were observed: (i) elevated core fucosylation (LcH) **(Figure 1A, 1E)** and (ii) a decrease in core 1/3 *O*-glycans compared to equine OA (AIA, MPL, MNA-G) **(Figure 1A)**.

**Figure 2:**
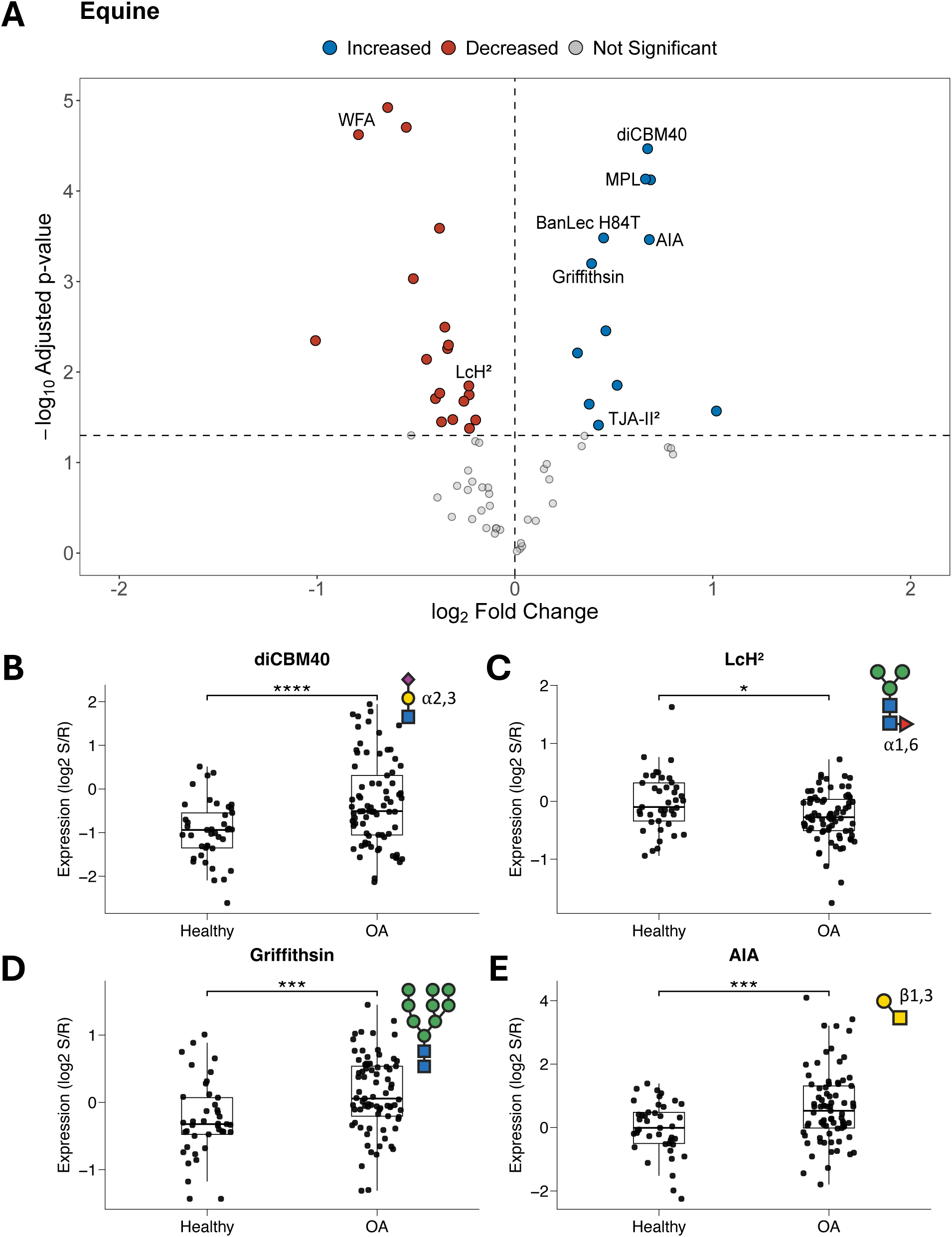
Glycomic alterations in equine synovial fluid associated with OA. (A) Volcano plot comparing lectin microarray signal intensities between OA and control horses. Blue and red dots represent lectins with significantly lower or higher signal intensities, respectively (FDR-adjusted *p* < 0.05). (B–E) Lectins recognizing α2,3-sialylation, core fucosylated glycans, high-mannose glycans, and core 1/3 O-glycans show altered binding in OA synovial fluid. *p* < 0.05, \**p* < 0.01, \*\**p* < 0.001, \*\*\**p* < 0.0001. Lectin labels are shown with abbreviated names; duplicated probes are distinguished by numeric superscripts (e.g., LcH¹, LcH²; anti-Leᵃ¹, anti-Leᵃ²). See Table S4 for full description.

**Figure 3:**
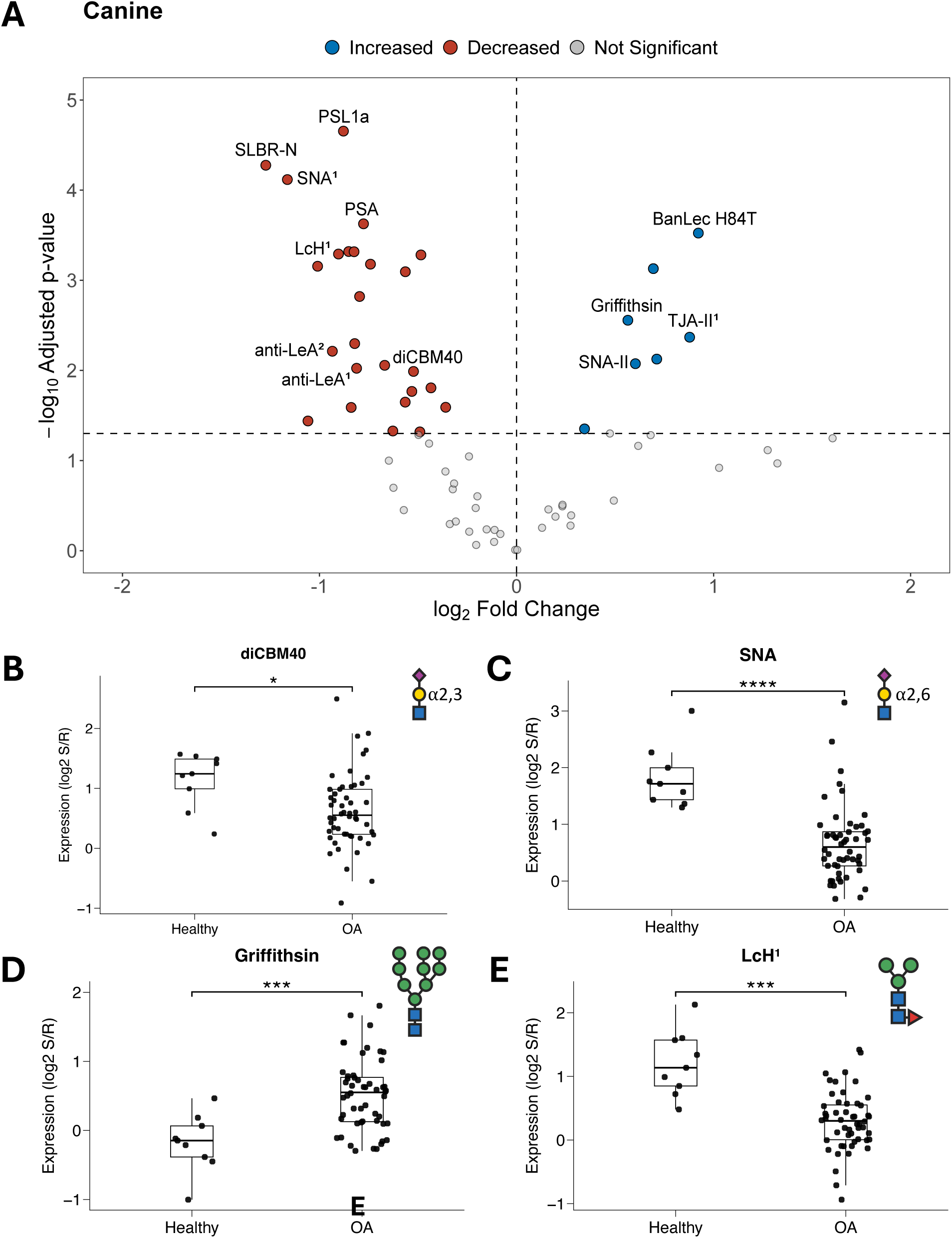
Glycomic alterations in canine synovial fluid associated with OA. (A) Volcano plot comparing lectin microarray signal intensities between OA and control dogs. Blue and red dots represent lectins with significantly lower or higher signal intensities, respectively (FDR-adjusted *p* < 0.05). (B–E) Lectins recognizing α2,3-sialylation, α2,6-sialylation, high-mannose glycans, and core fucosylated glycans show altered binding in OA synovial fluid. *p* < 0.05, \**p* < 0.01, \*\**p* < 0.001, \*\*\**p* < 0.0001. Lectin labels are shown with abbreviated names; duplicated probes are distinguished by numeric superscripts (e.g., LcH¹, LcH²; anti-Leᵃ¹, anti-Leᵃ²). See Table S4 for full description.

### Altered sialylation in canine and equine OA joints

Sialic acids are covalently linked monosaccharides typically located at the terminal positions of glycans in glycoconjugates, where they most often cap the glycan chains of glycoproteins and glycolipids. In humans, sialylation is typically added by the ST3 beta-galactoside alpha-2,3-sialyltransferases, ST6 beta-galactoside alpha-2,6-sialyltransferases, and ST8 alpha-N-acetyl-neuraminide alpha-2,8-sialyltransferases.

This categorization is based on the position of sialic acid addition (51–53). In canine and equine models of osteoarthritis, alterations in sialylation were observed in sera and synovial fluid, respectively (13, 54, 55). Given the crucial role of sialic acids in health and disease, we investigated their lectin-binding patterns in OA across species using lectin microarray analysis. Our analysis revealed a reduction in α2,6-sialylation in canine joints and a reduction in α2,3-sialylation in both canine and equine joints. These differences were mainly revealed by Sambucus nigra agglutinin (SNA-I), which binds α2,6-linked sialic acids (56), as well as by diCBM40 and the Maackia amurensis lectins MAL-I and MAL-II, which recognize distinct α2,3-sialylated structures. (57, 58). Notably, these findings are consistent with glycomic analyses of equine synovial lubricin, which showed a disease-associated shift from disialylated to monosialylated Core 1 *O*-glycans in OA joints, alongside a general reduction in overall sialylation (55).

### Changes in core fucosylation

N-glycan core fucosylation is a post-translational modification occurring in mammalian tissues. The sole glycosyltransferase, FUT8, transfers a fucose residue from GDP-Fuc to the innermost *N-*acetylglucosamine (GlcNAc) moiety of N-glycans and forms an α1,6-linkage (59, 60). Aberrant core fucosylation is an important glycosylation event because it alters protein conformation and function, contributes to immune dysregulation, and is strongly associated with inflammation, cancer progression, and metastasis (61–63).

Species-specific alterations in core fucosylation levels were evident across synovial fluid samples. In samples from equine and canine subjects, we detected diminished core fucosylation as indicated by significant reduction in *Lens culinaris* hemagglutinin, *Pisum sativum* agglutinin, and *Aleuria aurantia* lectin (AAL) binding **(Figures 2C, 3A, 3E)**.

These lectins preferentially recognize α1,6-linked core fucose (64, 65). In contrast, there is a strong increase in core fucose in human SF in OA **(Figure 1A, 1C)**.

### Changes in *O*-glycan structures

*O*-glycan biosynthesis begins with the addition of N-acetylgalactosamine (GalNAc) to serine or threonine residues, forming the Tn antigen. This structure can be extended by glycosyltransferases to produce distinct cores, with Core 1 (Galβ1-3GalNAc) and Core 3 (GlcNAcβ1-3GalNAc) among the most common, playing key roles in mucin architecture and joint lubrication (66–68). In synovial fluid, these glycans are largely carried by lubricin, encoded by PRG4, whose extensive O-glycosylation contributes to boundary lubrication, protease resistance, and immune modulation (67–70).

Our lectin microarray analysis revealed species-dependent changes in Core 1/3 O-glycan presentation in synovial fluid, likely reflecting differences in lubricin glycoforms. In equine OA joints, these glycans were increased, consistent with Noordwijk et al. (13) who reported elevated Core 1 O-glycans and higher lubricin levels using lectin microarray and metabolomic profiling. In canine SF, Wang et al. (71) reported increased lubricin and Core 1 O-glycans using PNA lectin and the anti-lubricin mAb MABT401 (9G3); however, our microarray detected only mannose and blood group antigens, with no increase in Core O-glycans. In human OA SF, Core 1/3 O-glycans detected by Artocarpus integrifolia agglutinin (AIA; Jacalin) and Macluria pomifera agglutinin (MPA)—which preferentially bind 3-substituted GalNAcα structures—were reduced **(Figure 2B)** (72–74). Interestingly, human lubricin levels have been reported as variable across studies (75, 76).

### High-mannose glycans are conserved in OA SF from humans, dogs and horses

High-mannose glycans are important in the biosynthesis of glycoproteins and quality control, ensuring proper protein folding and preventing the secretion of misfolded glycoproteins(77). Aberrant elevation of high-mannose structures has been associated with the pathogenesis of multiple malignancies, including colorectal and breast cancers (78, 79). Furthermore, high-mannose N-glycans have been associated with both pulmonary tissue injury and heightened systemic disease burden in the context of influenza virus infection (42, 80). Lectin microarray profiling revealed a conserved increase in high mannose glycans in synovial fluid from OA joints across all three species. This elevation was detected by increased binding of Griffithsin **(Figure 1-3D)** and H84T **(Figure 2-3F)**, which recognize mannose-rich N-glycan motifs (81–83), and illustrated in volcano plots **(Figure 1A-3A)**.

### Glycan changes were conserved between species despite different injuries

Despite differences in joints sampled and injury types, glycan alterations in canine and equine OA joints were more like each other than to those observed in humans. Equine synovial fluid samples were collected predominantly from carpal joints or other joints with osteochondral fragmentation, whereas human and canine samples were obtained from knee/stifle joints with anterior cruciate ligament/cranial cruciate ligament rupture. The stronger similarity between equine and canine profiles likely reflects differences in study design between veterinary patient and human patient samples. Whereas healthy and OA joints could be sampled independently from different individuals for the veterinary species, it is not ethically feasible to sample synovial fluid aspirates from human patients free of joint disease. Therefore, a limitation of the human samples is that both OA and contralateral control joints were paired, with both samples being obtained from the same individual in a subset of patients that gave permission for bilateral sampling. While idiopathic or age-related osteoarthritis is typically a bilateral disease, albeit affecting one joint more severely than another (84), the contralateral joint–especially for TKR patients–should not be truly considered a “healthy” joint.

Nevertheless, an increase in high mannose glycans was consistently observed across all three species, suggesting this may represent a core glycomic signature of OA. Previous studies have shown that glycan profiles can vary with age and sex, particularly in adult human populations. For example, Valerie et al. (85) reported age-related increases in glycan branching and decreases in galactosylation, while others have found sex-dependent differences in glycan abundance that become more pronounced after puberty (86, 87). Thus, we considered age and sex as potential confounding variables that could obscure or inflate observed OA-associated glycomic changes.

We performed covariate-adjusted linear regression models which included unadjusted, age-adjusted, and age-plus-sex–adjusted comparisons to assess whether demographic variables confounded OA associations (Data S2). Across the 55 lectins assessed, OA-associated differences were found in 24 lectins in dogs, 28 in horses, and 17 in humans (Table S5). Age-related associations were found for 6 lectins in dogs, 13 in horses, and 6 in humans, primarily reflecting variations in high-mannose, bisected GlcNAc, and terminal sialylation motifs. Sex-related effects were present in 4 lectins in horses and 6 in humans, enriched for fucosylated Lewis-type and sialylated epitopes. Similarly, our horse datasets lacked a balance of intact male subjects, thereby limiting the statistical power to fully resolve sex-related differences (Table 1). Due to these imbalances, observed sex effects in horses and dogs and age effects in humans must be interpreted cautiously. A major limitation of the human cohort is that contralateral “control” joints were obtained from the same individuals as the OA joints, meaning they cannot be considered truly healthy controls.

**Table 1:**
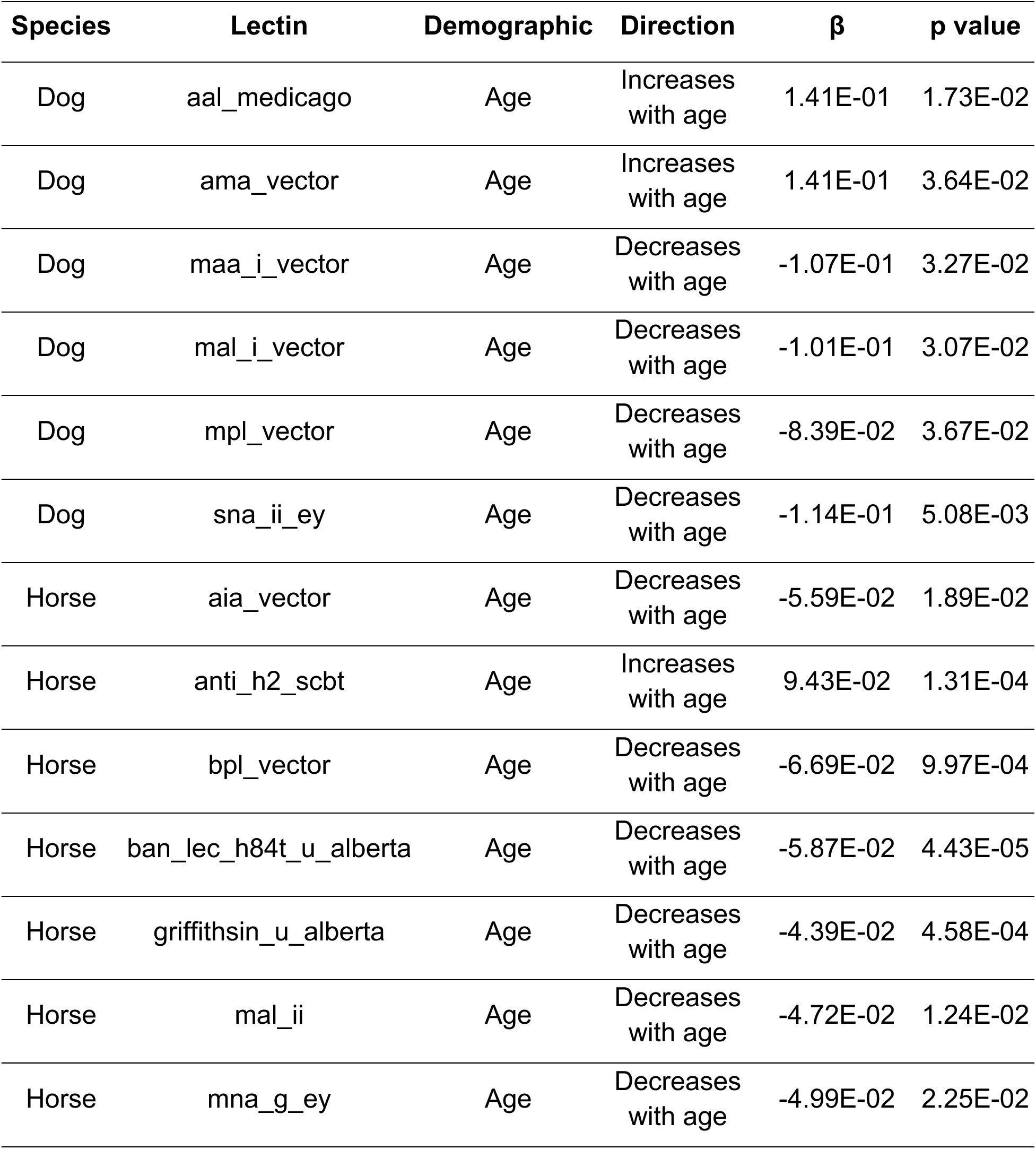
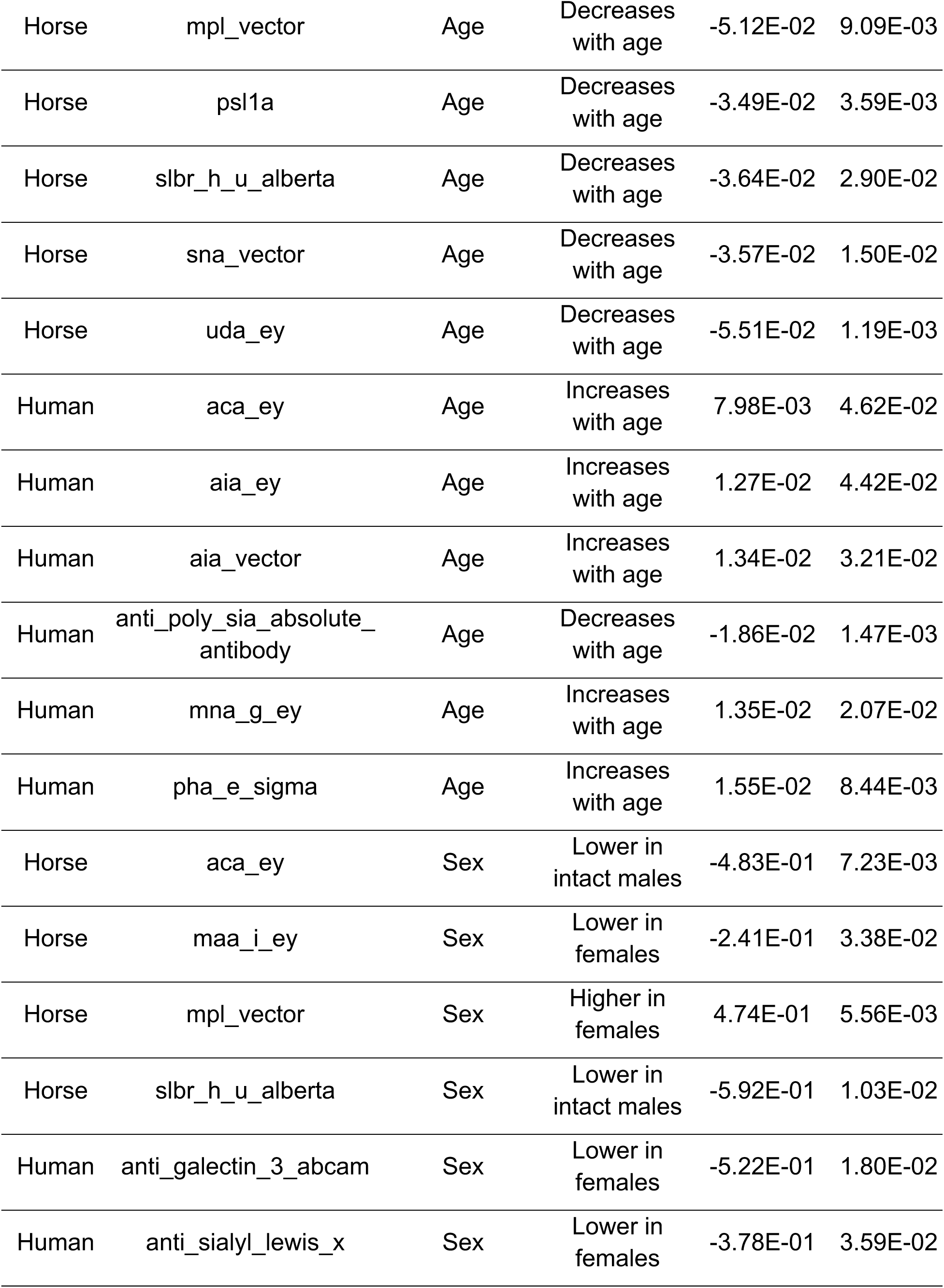
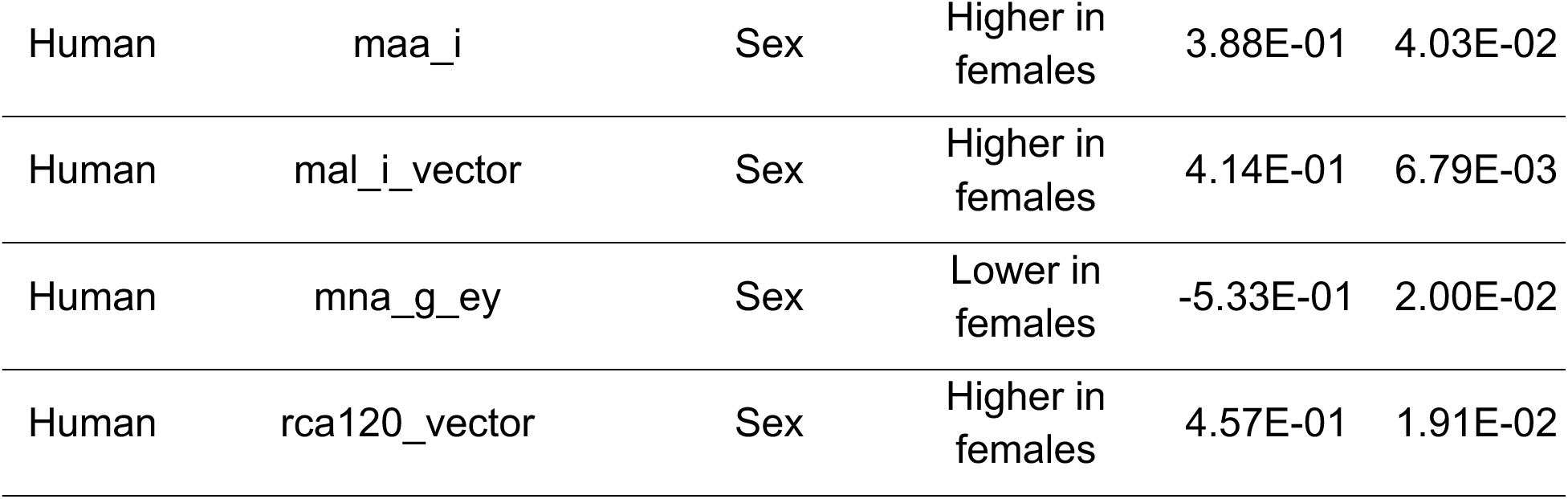
Age-and sex-associated lectins identified in canine, equine, and human synovial fluid samples. β = regression coefficient. p values calculated using values calculated using two-sided Wald t-tests on regression coefficients from ordinary least squares (OLS) linear models.

| Species | Lectin | Demographic | Direction | $\beta$ | p value |
| --- | --- | --- | --- | --- | --- |
| Dog | aal_medicago | Age | Increases with age | 1.41E-01 | 1.73E-02 |
| Dog | ama_vector | Age | Increases with age | 1.41E-01 | 3.64E-02 |
| Dog | maa_i_vector | Age | Decreases with age | -1.07E-01 | 3.27E-02 |
| Dog | mal_i_vector | Age | Decreases with age | -1.01E-01 | 3.07E-02 |
| Dog | mpl_vector | Age | Decreases with age | -8.39E-02 | 3.67E-02 |
| Dog | sna_ii_ey | Age | Decreases with age | -1.14E-01 | 5.08E-03 |
| Horse | aia_vector | Age | Decreases with age | -5.59E-02 | 1.89E-02 |
| Horse | anti_h2_scbt | Age | Increases with age | 9.43E-02 | 1.31E-04 |
| Horse | bpl_vector | Age | Decreases with age | -6.69E-02 | 9.97E-04 |
| Horse | ban_lec_h84t_u_alberta | Age | Decreases with age | -5.87E-02 | 4.43E-05 |
| Horse | griffithsin_u_alberta | Age | Decreases with age | -4.39E-02 | 4.58E-04 |
| Horse | mal_ii | Age | Decreases with age | -4.72E-02 | 1.24E-02 |
| Horse | mna_g_ey | Age | Decreases with age | -4.99E-02 | 2.25E-02 |
| Horse | mpl_vector | Age | Decreases with age | -5.12E-02 | 9.09E-03 |
| Horse | psl1a | Age | Decreases with age | -3.49E-02 | 3.59E-03 |
| Horse | slbr_h_u_alberta | Age | Decreases with age | -3.64E-02 | 2.90E-02 |
| Horse | sna_vector | Age | Decreases with age | -3.57E-02 | 1.50E-02 |
| Horse | uda_ey | Age | Decreases with age | -5.51E-02 | 1.19E-03 |
| Human | aca_ey | Age | Increases with age | 7.98E-03 | 4.62E-02 |
| Human | aia_ey | Age | Increases with age | 1.27E-02 | 4.42E-02 |
| Human | aia_vector | Age | Increases with age | 1.34E-02 | 3.21E-02 |
| Human | anti_poly_sia_absolute_antibody | Age | Decreases with age | -1.86E-02 | 1.47E-03 |
| Human | mna_g_ey | Age | Increases with age | 1.35E-02 | 2.07E-02 |
| Human | pha_e_sigma | Age | Increases with age | 1.55E-02 | 8.44E-03 |
| Horse | aca_ey | Sex | Lower in intact males | -4.83E-01 | 7.23E-03 |
| Horse | maa_i_ey | Sex | Lower in females | -2.41E-01 | 3.38E-02 |
| Horse | mpl_vector | Sex | Higher in females | 4.74E-01 | 5.56E-03 |
| Horse | slbr_h_u_alberta | Sex | Lower in intact males | -5.92E-01 | 1.03E-02 |
| Human | anti_galectin_3_abcam | Sex | Lower in females | -5.22E-01 | 1.80E-02 |
| Human | anti_sialyl_lewis_x | Sex | Lower in females | -3.78E-01 | 3.59E-02 |
| Human | maa_i | Sex | Higher in females | 3.88E-01 | 4.03E-02 |
| Human | mal_i_vector | Sex | Higher in females | 4.14E-01 | 6.79E-03 |
| Human | mna_g_ey | Sex | Lower in females | -5.33E-01 | 2.00E-02 |
| Human | rca120_vector | Sex | Higher in females | 4.57E-01 | 1.91E-02 |

Despite observing these demographic effects, the strongest OA markers including high-mannose, core O-glycans, and terminal sialylation epitopes remained significant across all species, with overlapping confidence intervals between unadjusted and adjusted regression models. In short, while we found that age and sex influence certain glycan motifs, they do not obscure the primary OA-associated glycomic patterns. Rather, age and sex act as modulators that add biological context to the OA signatures, and the observed changes are disease-driven but demographically nuanced. Since age emerged as a recurrent covariate across species, we next examined whether OA severity was directly associated with age. This analysis was performed only in the equine cohort, because (i) OA grade data were only available for horses, and (ii) contralateral sampling in humans precludes reliable assignment of true “healthy” grades. Jittered scatterplots with regression smoothing revealed a weak positive trend between age and OA grade **(Figure S2)**. Pearson’s correlation supported this trend (r = 0.16, 95% CI: –0.02 to 0.33, p = 0.085). These findings indicate that while age possibly contributes to OA severity in horses, it does not fully account for the glycomic alterations observed.

### CatBoost machine learning algorithms successfully separated OA from healthy individuals

Finally, we evaluated the discriminative power of glycan epitopes from our microarray for OA versus healthy control classification using the PyCaret library (version 3.3.2) (88). A binary classification framework was applied to distinguish between controls (0) and OA joints (1) based on lectin binding profiles. The combined dataset (Data S1) consisted of 60 control and 166 OA samples, each characterized by 68 numerical features, and was randomly split into a 70:30 training set:validation set. We employed a stratified ten-fold cross-validation scheme (stratified by condition) to maintain consistent class proportions across folds.

To address class imbalance, we applied the Synthetic Minority Over-Sampling Technique (SMOTE) to the training set using the scikit imbalanced-learn library (version 1.4.2.). The performance of multiple machine learning models (accuracy, area under curve[AUC], recall, precision, F1, kappa, and Matthew Correlation Coefficient [MCC]) in the training cohort is shown in Table S6. While the Extra Trees and Light Gradient Boosting Machine achieved better overall performance, CatBoost offered comparable performance but with several advantages. Notably, its ordered boosting strategy and regularization mechanisms reduce overfitting (89, 90), which is particularly beneficial for small, imbalanced datasets. Given these interpretability and generalizability benefits and with minimal tradeoff in performance, CatBoost was selected as the optimal model. CatBoost demonstrated strong mean cross-validation performance using default hyperparameters (Table 2). Hyperparameter tuning did not yield consistent improvements (data not shown); therefore, we retained the untuned CatBoost model.

**Table 2:** Performance of the CatBoost model distinguishing control (0) versus OA (1) samples using stratified 10-fold cross-validation. Values are means ± standard deviation (SD) across 10 iterations. Metrics: AUC, area under the receiver operating characteristic curve; MCC, Matthew’s correlation coefficient; F1, F1 score; Kappa, Cohen’s kappa.

|  | Accuracy | AUC | Recall | Precision | F1 | Kappa | MCC |
| --- | --- | --- | --- | --- | --- | --- | --- |
| Mean | 0.91 | 0.98 | 0.86 | 0.97 | 0.90 | 0.82 | 0.83 |
| Std Dev | 0.05 | 0.02 | 0.12 | 0.06 | 0.06 | 0.09 | 0.08 |

Given its high cross-validation scores and architectural advantages, CatBoost appeared well-suited to capture the intricate glycan–epitope interactions underlying OA pathophysiology across species.

The binary classification model demonstrated strong classification performance on the holdout set, as shown in **Figure 4C**. It correctly identified 100% of OA cases (17 out of 17 samples), achieving perfect recall. Specificity was moderate at 50%, with 3 out of 6 healthy controls misclassified as OA, but the model achieved an overall accuracy of 86.96% (20/23). This classification pattern reflects a deliberate bias toward maximizing sensitivity, as the model excelled in identifying OA cases while demonstrating only modest performance for healthy controls. It should also be noted that the balance between sensitivity and specificity is not fixed, but can be influenced by adjusting model thresholds, as the optimal balance may differ depending on the clinical or research context.

**Figure 4:**
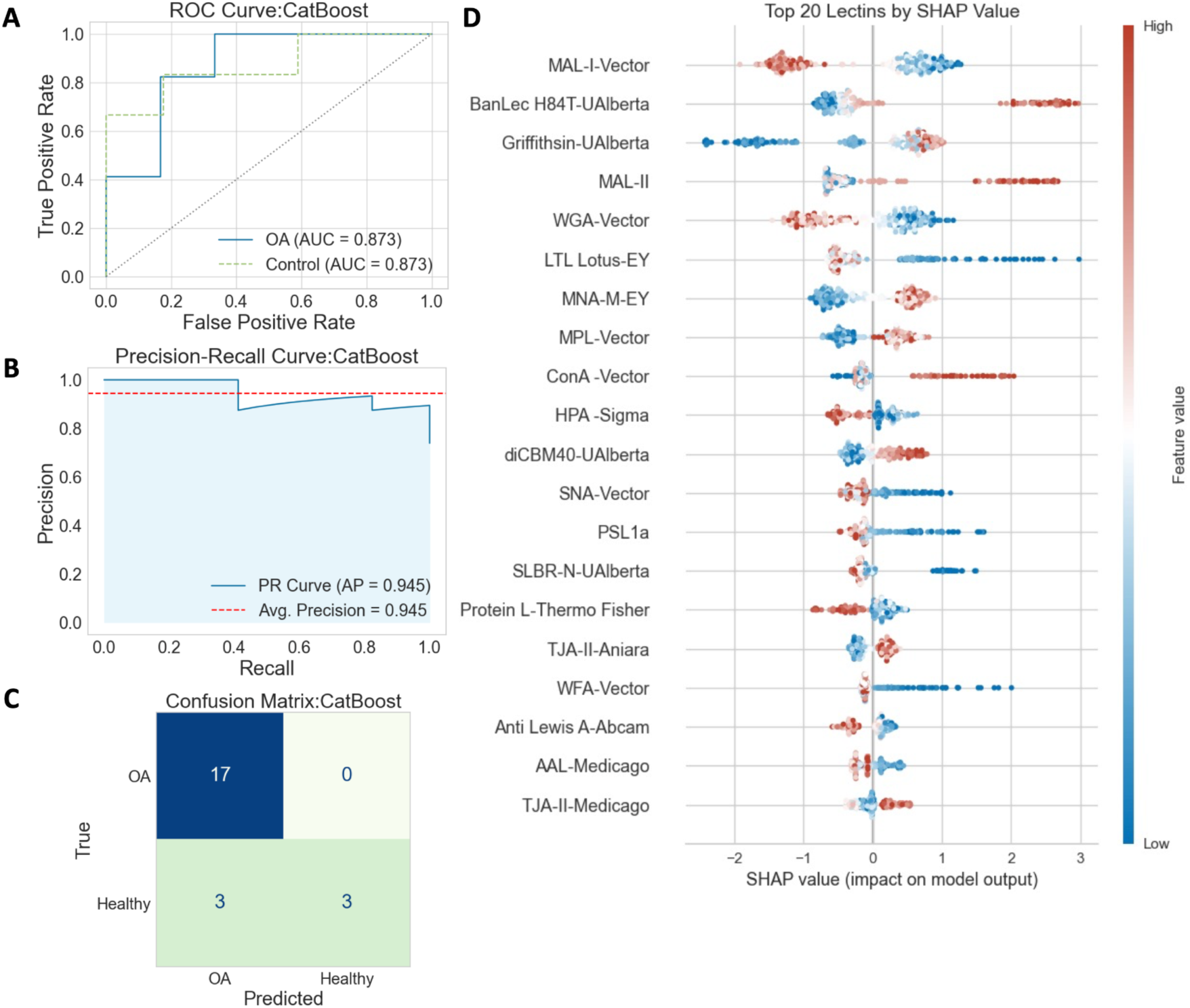
Machine learning classification of OA using CatBoost models across species. (A) Receiver operating characteristic (ROC) curve of the CatBoost classifier distinguishing OA from control samples. (B) Precision-recall (PR) curve for the same model, with the solid line representing the binary PR curve and the dashed line indicating average precision. (C) Confusion matrix showing CatBoost predictions versus true class labels in the holdout set. (D) SHAP analysis of feature contributions to OA classification. The top 20 lectins ranked by mean absolute SHAP value are shown, with beeswarm plots illustrating the distribution of SHAP values across samples. Dot colors indicate relative lectin signal intensity, scaled by percentile within the feature’s distribution.

To further assess model discrimination on the holdout set, we evaluated the receiver operating characteristic (ROC) and precision–recall (PR) curves. CatBoost achieved an area under the ROC curve (AUC) of 0.873 for both OA and healthy classes **(Figure 4B)**, indicating balanced discriminative ability. The precision–recall curve yielded an average precision (AP) score of 0.945 **(Figure 4A)**, highlighting the model’s robustness in handling class imbalance and reinforcing its suitability for OA classification. Moreover, the model’s performance was further evaluated by analyzing the learning curve **(Figure S3A)**, which illustrates the progression of AUC scores for both the calibration and validation sets as the number of training instances (x-axis) increases. The AUC of the model is represented by a green line with 10 markers, each marker corresponding to a calculation performed using 10-fold cross-validation (CV). This AUC curve is accompanied by a shaded green region, which represents the range between the minimum and maximum AUC values obtained by CatBoost during CV. Alongside, the blue line represents the training score (AUC on the training set), which illustrates the model’s performance under ideal conditions and serves as a comparative benchmark. It can be observed that the training AUC quickly reaches 1.0 with fewer than 50 training samples, while the cross-validation AUC remains below 0.9 with the same number of training instances. Similarly, the validation curve **(Figure S3B)**, where the x-axis denotes the maximum tree depth, demonstrates that the AUC score begins above 0.96 at a tree depth of 2 and rapidly rises to above 0.99 as the maximum tree level reaches 10. The consistently high AUC scores for both the training and validation sets indicate that this model has sufficient capacity to fit the training data (training AUC = 1.0), while the steadily improving validation AUC demonstrates that the model’s generalization ability improves as more data are included and as the model’s capacity (tree depth) increases.

To obtain additional understanding into the machine learning model’s results, we used SHAP (SHapley Additive Explanations) analysis. We applied it on the training data to ensure methodological rigor and limit information leaking. This approach revealed consistent feature importance patterns that support the robustness of the selected classifier. SHAP visualized the contribution of each specific lectin and its associated glycan epitopes, with SHAP values indicating both the direction and magnitude of each feature’s impact on the model. SHAP importance values quantify the average marginal contribution of each feature, in our case, individual lectins—to the model’s prediction, across all possible feature combinations (91, 92). Unlike traditional feature importance rankings, SHAP provides case-level interpretability by showing how specific glycan-binding profiles influence individual classification outcomes. In the summary plot **(Figure 4D)**, high (red) and low (blue) lectin signal intensities differentially push predictions toward OA or healthy, offering a nuanced understanding of how glycan epitopes contribute to disease classification beyond what permutation importance alone can reveal.

Expanding on these findings, SHAP values illustrate how glycan-binding signals from specific lectins shape OA classification decisions. BanLec and MAL-II lectins show broad SHAP value distributions, with higher binding intensities (red) corresponding to positive SHAP values, indicating that strong binding strongly predicts OA. In contrast, Griffithsin displays extreme SHAP values at low binding intensities (blue), showing that weak binding has a strong negative impact on the model. Overall, these SHAP profiles suggest that BanLec, MAL-II, and Griffithsin are high-impact predictors of OA. Conversely, SNA-I and MAL-I also exhibit broad SHAP distributions, but in their case, low binding intensities (blue) are associated with positive SHAP contributions, suggesting that reduced recognition by these lectins is likewise characteristic of the OA class. In contrast, WFA and Anti-Sialyl Lewis X display narrower SHAP distributions, indicating lower overall model impact. However, both show a trend where lower binding values contribute positively to OA prediction, albeit with less influence compared to the top-ranked features. In summary, our SHAP analysis not only enhances our model’s explainability but also identifies key glycan features for further biological investigation.

## Discussion

The diagnosis of early osteoarthritis is often delayed because of the poor sensitivity of currently available diagnostic tests, which rely on patient history, physical exam findings, and imaging findings which often lag behind symptomology. Synovial fluid, by virtue of its direct contact with cartilage and synovium, represents a biologically informative substrate for OA biomarker discovery. Our findings show that glycomic profiling of synovial fluid captures the glycosylation changes associated with OA, suggesting that glycan biomarkers offer an informative and accessible diagnostic approach that may be more sensitive than current methods. These glycan biomarkers include high-mannose, sialylation, core-1 *O*-glycans, and core fucose.

In OA, a consistent reduction in α2,6-linked sialylation emerges as a key molecular signature with potential functional consequences. Transcriptomic and glycomic profiling of arthritic synovial fibroblasts (ASFs) revealed downregulation of ST6GAL1 and reduced SNA lectin binding, supporting a shift away from α2,6-sialylation toward a glycome enriched in terminal galactose and α2,3-sialylation (93). This desialylated state increases galactose exposure, enhancing galectin-3 binding (94) and subsequent induction of IL-6 and CCL2, inflammatory cytokines that further suppress ST6GAL1 expression, creating a self-reinforcing inflammatory loop (95). Critically, α2,6-sialylation functions as a molecular checkpoint for Siglec-5 signaling; its loss impairs both the expression and engagement of this inhibitory receptor, removing a key brake on TLR4-driven inflammation. The linkage-specific remodeling also decreases engagement of other anti-inflammatory Siglecs while favoring recognition by pro-inflammatory receptors like Siglec-1 (sialoadhesin), which is upregulated on activated macrophages and promotes T cell infiltration (17, 96). This sialylation imbalance also affects secreted glycoproteins such as lubricin, a key joint lubricant with emerging immunomodulatory roles. In equine OA, lubricin levels are increased but exhibit elevated non-sialylated *O*-glycans in its mucin domain, as detected by PNA lectin and 9G3 antibody binding (71). Given that lubricin interacts with CD44 and Toll-like receptors 2 and 4 to inhibit synovial inflammation (97, 98), a reduction in terminal sialylation may impair its ability to engage these receptors and maintain immunological quiescence. Together, these findings suggest that the pro-inflammatory shift in sialylation—both at the articular cartilage surface and on secreted glycoproteins like lubricin—may represent a mechanistic driver of sustained joint inflammation and tissue degradation in OA.

There is emerging evidence highlighting core fucosylation as an essential modulator of cartilage homeostasis and glycan-mediated signaling. Fut8 conditional knockouts in mouse chondrocytes result in diminished TGF-β signaling, elevated MMP13 expression, and accelerated cartilage degradation, which are all indicative of early and progressive OA (99). This is consistent across both injury-induced and age-related models, implicating core fucosylation as a structural and signaling checkpoint in OA (99). In particular, Fut8(-/-) mice were reported to exhibit suppressed phosphorylation of Smad and increased expression of MMPs (matrix metalloproteinases) due to decreased binding of TGF-β ligand attributed to the lack of core fucose addition to the TGF-β type II receptor (59, 100).

Building on this, recent experimental studies by Homan et al. (99) showed that mannosidase-induced trimming of high-mannose N-glycans—specifically the reduction of Man8–9GlcNAc2 and accumulation of smaller oligomannose species such as Man5– 6GlcNAc2—promoted reversible OA-like changes, including proteoglycan loss and increased nitric oxide release from rabbit and mouse articular cartilage. These trimmed glycans created substrates for FUT8, thereby facilitating core fucosylation as a compensatory “repair” mechanism that helps stabilize the extracellular matrix and preserve chondrocyte homeostasis. In the absence of FUT8, however, mannosidase treatment exacerbated degeneration, marked by suppressed *Tgfb* expression and upregulation of *Mmp13*. Together, these data establish high-mannose processing and subsequent core fucosylation as an interdependent checkpoint in glyco-mediated cartilage preservation.

Our cross-species comparisons revealed divergent patterns of core fucosylation across species: fucosylation was increased in human OA synovial fluid but decreased in both canine and equine OA samples compared to healthy controls. As such, understanding the contribution of altered fucosylation to OA pathophysiology will require placing these differences in the context of upstream glycan processing events.

Previous studies on human and mouse cartilage samples have reported reductions in high-mannose in OA due to a decrease in ConA binding and immunohistochemical staining, respectively (38, 101). These cartilage findings contrast with our lectin microarray findings across dog, horse, and human synovial fluid samples. However, it is important to note that cross-comparison analyses of glycan microarray revealed a broader binding profile for ConA that includes biantennary and complex N-glycans, and ConA binding can be inhibited by α1,2-or α1,3-linked fucosylation at glycan termini (74). In OA, increased terminal fucosylation or branching may reduce ConA signals, even in the presence of high-mannose glycans. Conversely, BanLec and Griffithsin are more selective for high-mannose structures (especially Man8–9) (81, 102, 103), likely providing a clearer picture of high-mannose abundance. Furthermore, as previously mentioned, unlike cartilage-based studies, we examined synovial fluid (SF), a joint-derived biofluid that provides a highly localized molecular snapshot of OA pathophysiology due to its direct contact with articular cartilage, synovium, ligaments, and other affected tissues. This spatial proximity enables SF to reflect early and dynamic biochemical changes that may not be fully represented in cartilage analyses (Mickiewicz et al., 2015). Our cross-species results therefore complement existing cartilage studies and suggest that high-mannose accumulation may be a conserved feature of OA across multiple joints.

Interestingly, while high-mannose glycans were consistently elevated across all three species in our study, the expected compensatory increase in core fucosylation was observed only in human OA synovial fluid, but not in dogs or horses, where it was markedly reduced. These findings raise the possibility that, although high-mannose accumulation is a conserved feature of OA, the capacity to initiate a core fucosylation– based repair response may differ by species, potentially reflecting variation in FUT8 activity or the timing of disease progression. Supporting this, species-specific differences in glycosyltransferase activity may help explain the divergence. For example, ST3GAL2 is expressed in human but not mouse bone marrow and shows different substrate preferences across species (104, 105). Similarly, variation in FUT8 expression or enzyme efficiency in dogs and horses could account for the lack of compensatory core fucosylation despite high-mannose accumulation.

Because age and sex are well-established modulators of glycosylation, their potential to influence glycan signals in synovial fluid requires close examination, especially when interpreting osteoarthritis-associated glycomic changes in microarray analyses. Our dataset revealed species-specific variations in glycan motifs linked with high-mannose, core-*O*-glycans, and α2,6-sialylation. For instance, in horses, 13 lectins demonstrated age-associated shifts and 4 were influenced by sex, primarily within female and intact male subgroups. In humans, six lectins were age-associated and another six with the female sex. No significant sex effects were detected in dogs, this is likely due to sample imbalance, wherein all canine healthy controls were MI and canine OA were mainly FS and MN, thereby limiting the detection of sex differences.

A majority of OA-associated glycan signals remained significant after adjusting for age and sex, with effect directions unchanged. This suggests that these factors modify baseline abundance but do not drive the OA-related changes. Moreover, it aligns with prior human and animal studies demonstrating age-related increases in glycan branching and sialylation, and sex-dependent shifts in fucosylation (86, 106, 107). As a result, while age and sex influence the synovial glycome, core OA feature such as elevated high-mannose structures and decreased α2,6-sialylation are conserved across species The absence of significant sex associations in dogs is likely due to demographic imbalance instead of a true lack of sex-based glycomic differences. The OA group was predominantly composed of female spayed (FS) and male neutered (MN) individuals, which reflects common veterinary demographics. The lack of sex category overlap between the OA and control groups reduces the statistical contrast required to identify sex-related differences. Similarly, our equine cohort included relatively few intact males, with the majority of males being geldings, also reflecting common veterinary demographics. These demographic imbalances underscore the need for representative sampling—not as a constraint on glycomic investigations, but as an important factor in utilizing the full diagnostic potential of glycan-based biomarkers in OA research.

Machine learning techniques are helpful in identifying elusive patterns that are difficult to detect using conventional statistical methods and to test their predictive performance at the individual level (108–110). This study investigated machine learning approaches that utilizes glycome epitope signatures to classify OA and control subjects. As previous studies have demonstrated the effectiveness of CatBoost for classification tasks (111–113), our CatBoost model uncovered a subset of glycan epitopes with strong diagnostic potential for osteoarthritis by capturing conserved glycosylation features across species. Moreover, SHAP-based interpretation revealed that high mannose–binding lectins BanLec (mean SHAP value: 0.84) and Griffithsin (mean SHAP value: 0.82), as well as sialic acid–binding lectins MAL-I for α2,3 (mean SHAP value: 0.89) and SNA for α2,6 (mean SHAP value: 0.26), were among the top contributors to the model with higher mean SHAP values indicating greater feature importance. These same lectins also highlighted epitopes that were consistently enriched or depleted across human, canine, and equine OA samples, reinforcing their biological and diagnostic relevance. By offering individualized explanations for these predictions, this machine learning–based glycan profiling approach also holds promise for patient stratification, personalized treatment planning, and informed clinical decision-making.

While our study contributes a multimodal approach, several of its limitations warrant discussion. A key limitation of cross-species glycomic analysis is the lack of standardized nomenclature and incomplete structural annotation of glycans across organisms (114–117). Unlike proteins, glycans are not template-directed by an organism’s genome, and their biosynthesis depends on species-specific glycosyltransferase expression, substrate availability, and cellular context (118–120). Therefore, structurally similar glycan motifs may not be functionally equivalent across species. Although we found widely expressed epitopes like high mannose, which is known to interact with pro-inflammatory lectins like MBL2 and DC-SIGN (121), as well as core fucose, and α2,6-sialylation that seem to promote related biological processes, linkage-specific structural validation and enzyme-level expression data are necessary to demonstrate their functional equivalence.

Furthermore, dog and horse synovial fluid samples were obtained from separate individuals, whereas human samples were paired from the same patient. Contralateral sampling offers a built-in control for individual-level variation; however, it may underestimate glycosylation differences due to shared systemic influences or compensatory loading in the contralateral limb (122). Subclinical pathologies or early-stage OA in the control contralateral joint may partially conceal glycomic alterations, possibly explaining why glycan epitope differences exist between human and other large animal studies. Particularly in human patients undergoing TKR, it is common for the contralateral knee to have sufficiently advanced OA to require subsequent joint replacement. Even in patients who only have symptoms in one knee at the time of TKR, approximately 25% of patients undergo contralateral TKR within several years (123).

Finally, the presence of unbalanced data and a relatively small sample size, particularly in the healthy control group, may limit the generalizability of our findings. In this case, the class imbalance may have contributed to more accurate classification of osteoarthritis cases, as the model was trained on a larger number of OA samples than controls. To address this, we applied stratified data splitting, SMOTE, internal tenfold cross-validation, and comprehensive performance metric reporting. Another important limitation is the absence of testing the model on an external cohort, which restricts our ability to evaluate its generalizability across independent datasets. The limited sample size also prevented robust sex-based machine learning analyses, despite known differences in OA presentation, progression, and treatment response between sexes (124–127). Although the model performed well within our study cohort, future validation using larger and more balanced datasets is essential for assessing its generalizability and identifying any potential performance biases. Further studies should also explore whether incorporating additional biomarkers or covariates could improve the model’s predictive accuracy. As a logical consequence of our preliminary work, however, the study provides initial insights and highlights the potential of glycan-based profiling to capture molecular changes at the subject level, which may ultimately be applicable to early-stage or preclinical OA in future studies.

One Health based approaches are gaining momentum for optimizing biomarker discovery and informing translational therapeutic development (128–131). Here, multispecies glycomics analysis revealed high-mannose as a conserved glycan signature that was consistently upregulated in osteoarthritis across humans, dogs, and horses. In addition to this conserved feature, several other glycan epitopes displayed species-specific patterns **(Figure 5)**. Future studies should aim to uncover the mechanisms driving high-mannose accumulation in OA synovial fluid and determine whether this glycan alteration simply reflects downstream tissue remodeling or plays an active regulatory role in joint degeneration. Similarly, the divergent regulation of core fucosylation, sialylation and *O*-glycans across species raises important questions about the underlying cellular and enzymatic factors shaping the synovial glycome during disease progression.

**Figure 5:**
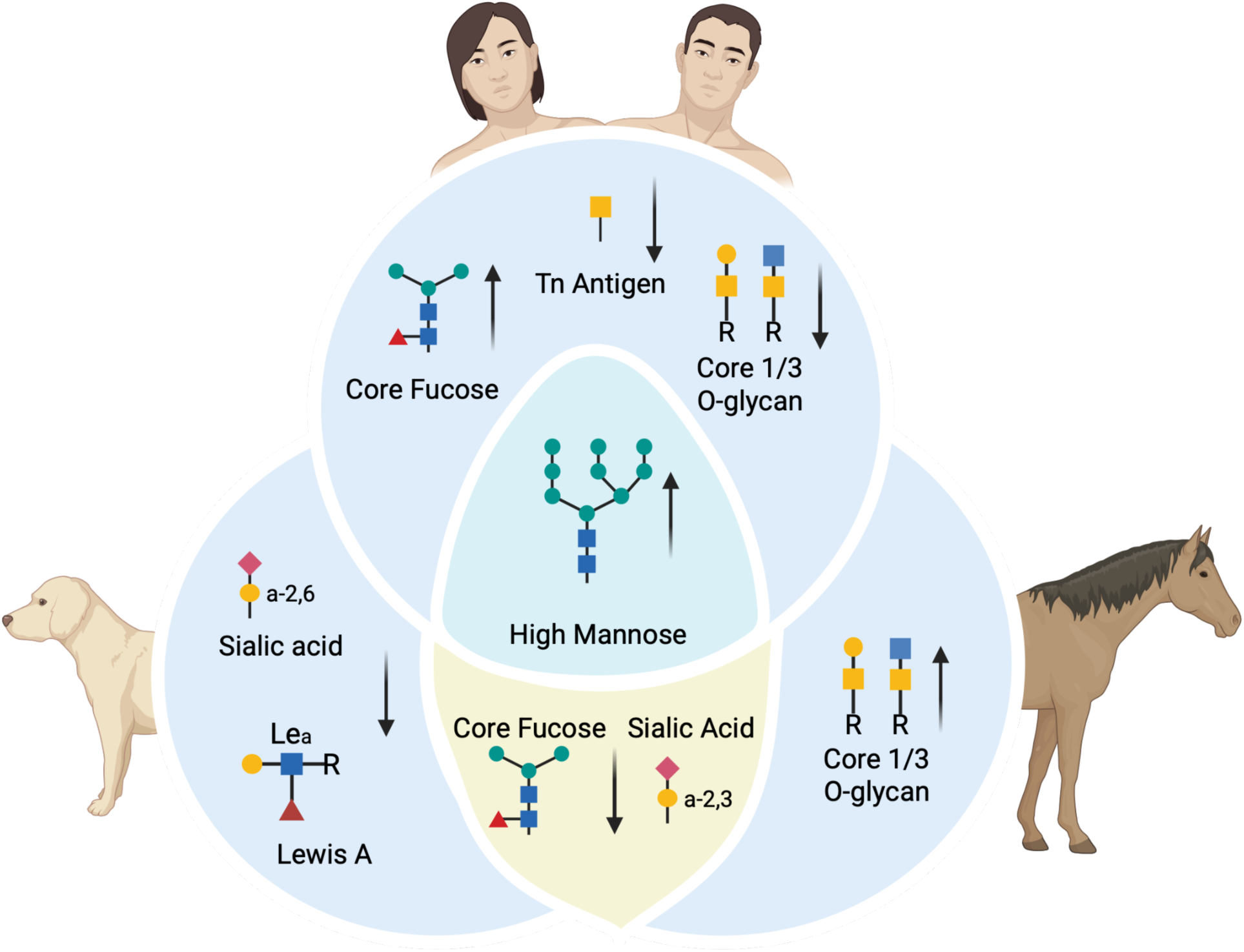
Comparative glycomic alterations in osteoarthritis (OA) across human, canine, and equine synovial fluid. Cross-species analysis highlights both conserved and species-specific glycan changes. High-mannose N-glycans are consistently enriched across all three species, representing a shared OA signature. Human OA is characterized by increased core fucosylation and decreased core 1/3 *O*-glycans. Canine OA exhibits reduced α2,6-sialylation, while equine OA shows increased core 1/3 *O*-glycans. Both equine and canine OA share decreases in core fucosylation and α2,3-sialylation. Created with BioRender.

A key contribution of this work is by combining glycomic profiling with advanced machine learning techniques to analyze the glycomic landscape of osteoarthritis across species. By using a unified analytical framework, we surpassed typical single-species or single-modality investigations, providing a more comprehensive understanding of glycan alterations involved in OA pathogenesis. While the machine learning models were implemented using established platforms such as PyCaret, our analytical pipeline can be customized, tested, and expanded for future studies with larger and more balanced datasets. This data-driven methodology highlights the translational potential of glycan-based biomarkers and lays the groundwork for future research into OA-related disease processes and diagnostic tools.

## Methods

### Experimental Design

#### Human Population

SF was collected from patients with PTOA of the knee secondary to anterior cruciate ligament (ACL) injury (n = 25) or from patients undergoing total knee arthroplasty (TKA) for idiopathic osteoarthritis (n = 11) presenting to the Hospital for Special Surgery (HSS). In a subset of patients, contralateral “healthy” knee joints were also sampled (n = 10 for ACL, n = 6 for TKA). All procedures were conducted with written informed consent, and study protocols were approved by the HSS Institutional Review Board (#2018-0490) in accordance with the Declaration of Helsinki. Patient data were anonymized, and demographic details are provided in Table S1.

#### Equine Population

Synovial fluid (SF) was obtained from horses with post-traumatic osteoarthritis (PTOA) involving the carpus (most commonly the middle carpal joint [MCJ] or antebrachiocarpal joint [ACJ]) as well as other joints including the fetlock (metacarpo-/metatarsophalangeal joint [MCPJ/MTPJ]), tarsus, and stifle / knee (n = 77), and from horses with healthy joints serving as controls (n = 41). Horses with PTOA presented to the Cornell University Hospital for Animals (CUHA) for arthroscopic treatment of carpal osteochondral fragmentation or subchondral bone/cartilage impact injury. Healthy carpal SF samples were obtained from horses free of musculoskeletal disease based on the absence of clinical lameness and either gross or arthroscopic findings and were collected within 30 minutes of euthanasia. SF was collected using aseptic technique immediately prior to joint distention for arthroscopy or at post-mortem collection. All sampling was performed with informed owner consent where applicable, and research procedures were approved by the Cornell University Institutional Animal Care and Use Committee (#2005-0015, #2018-0024). Demographic details for the equine cohort are provided in Table S2.

#### Canine Population

SF was collected from dogs with PTOA of the stifle / knee joints (n = 53) presenting to the Cornell University Hospital for Animals (CUHA) for surgical treatment of rupture of the cranial cruciate ligament (CCL). Healthy SF samples (n = 16) were obtained from dogs free of musculoskeletal disease based on the absence of clinical lameness and either gross or arthroscopic findings. All animal sampling was conducted under approved institutional protocols (#2005-0015, #2018-0024) with owner consent where applicable. Demographic details for the canine cohort are provided in Table S3.

#### Lectin Microarray Analysis

SF samples were processed for lectin microarray analysis as previously described in Noordwijk et al. (13). A schematic overview of the workflow is shown in **Figure 6**. In brief, synovial fluid was digested with hyaluronidase at 37° C for 1 hour. Samples (25 µg of protein) were labeled with NHS-activated Alexa Fluor 555 and a pooled reference sample was labeled with Alexa Fluor 647. Equal amounts of sample and reference (5µg) were hybridized on each array as previously described (132). Probes whose SNR \5 for^90% of samples were excluded. For the remaining probes, the intensity of each probe in each fluorescence channel was normalized to the median of the intensities of all probes. Lectins were annotated using Bojar et al. (74) and other literature (48–50, 83). The additional experimental information can be found in the supplementary file. A full breakdown of the lectin microarray workflow is also available in accordance with MIRAGE standards (Table S8)(47).

**Figure 6:**
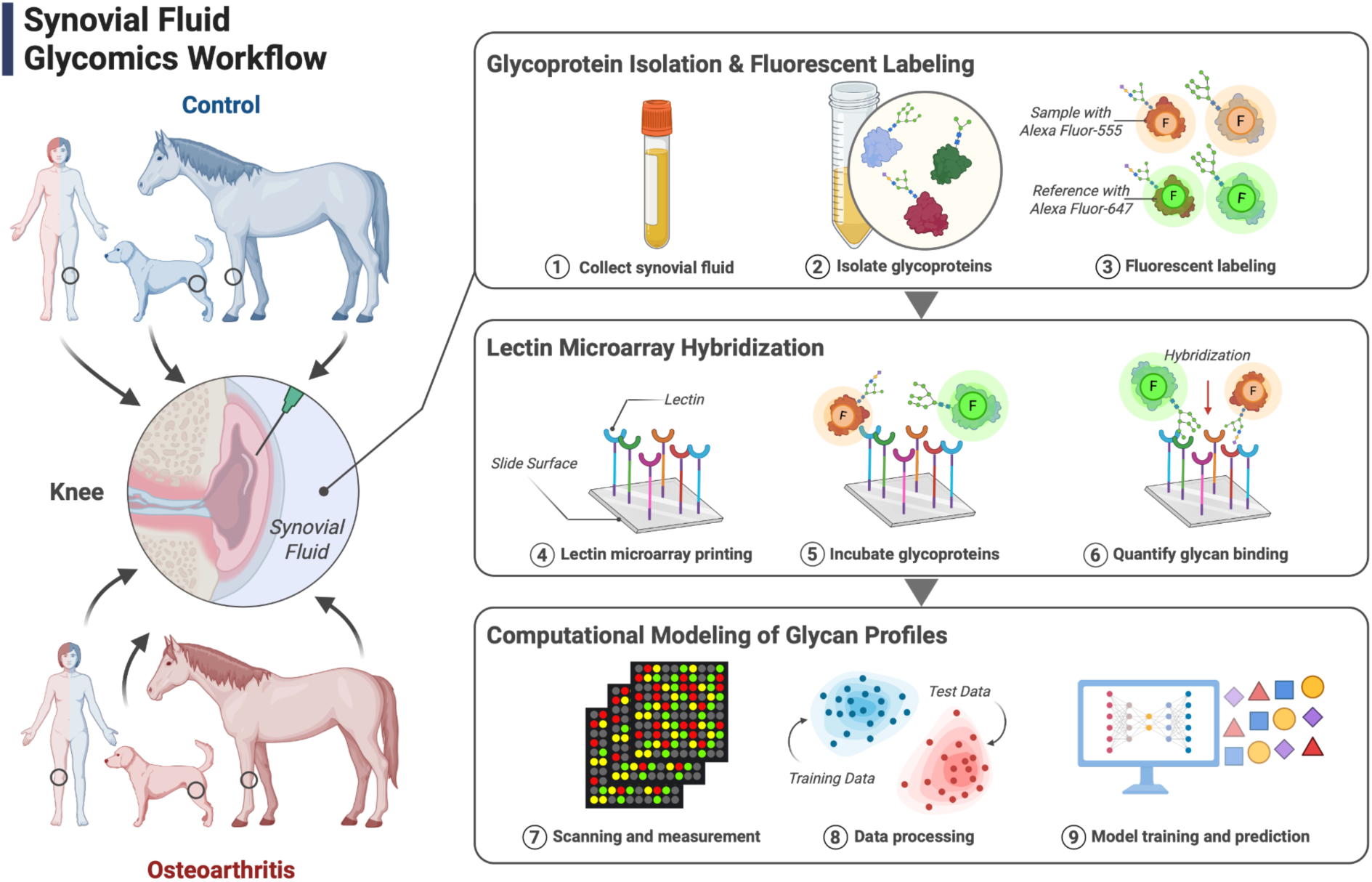
Workflow for lectin microarray analysis of synovial fluid glycoproteins. Schematic of the experimental pipeline: (1) synovial fluid collection, (2) glycoprotein isolation, (3) fluorescent labeling, (4–5) hybridization to lectin microarrays, (6) quantification to generate glycan-binding profiles, and (7–9) computational modeling and data analysis to identify OA-associated glycan signatures. See Experimental Procedures for full details. Created with BioRender.

#### Machine Learning Pipeline

We constructed a machine learning pipeline using Python (version 3.10.11) and PyCaret (version 3.0.0). The dataset underwent preprocessing, employing the Synthetic Minority Over-sampling Technique (SMOTE) to address class imbalance. The selected features were normalized using the MinMaxScaler from scikit-learn (version 1.4.2).

Model development incorporated features such as Species, Condition, and the binding signal intensities of 52 glycan epitope-specific lectins, 13 antibodies targeting glycan epitopes and immune-related proteins, and 3 Fc-binding proteins. Following a 10% holdout set, a training set and test set ratio of 7:3 was used, and stratified ten-fold cross-validation was performed to assess the model’s performance and robustness. We compared the performance of several machine learning models to identify the optimal model for classifying multi-species OA. This included Gradient Boosting (GBC), Light Gradient Boosting (LightGBM), Support vector machines (SVM), Extreme Gradient Boosting (XGBoost), Random Forest (RF), categorical boosting (CatBoost), Adaptive boosting (Ada), Extra Trees (ET), Logistic Regression (LR), and K Neighbors (KNN).

Their predictive performance was assessed using metrics such as F1, Kappa, MCC, Accuracy, AUC, Recall and Precision. Model hyperparameters were further tuned using grid search (default PyCaret function) on each model type to yield a higher AUC that was calibrated to the outcome across probability scores. The best model (highest AUC on the validation fold) was then set as the final model. While the best model was found to be ET and LightGBM, we chose CatBoost because of its ordered boosting strategy and regularization mechanisms to reduce overfitting which is particularly beneficial for small, imbalanced datasets. We reported ROC curves and AUC values for each fold and calculated the average AUC (where 1 = best classifier, 0.5 = random classifier. We created a PR (precision-recall) curve for each time point with an AP (average precision) value. Additionally, learning and validation curves were generated to assess the model’s ability to learn underlying trends in the data effectively and can generalize this learning to new data.

Subsequently, the Shapley Additive exPlanation (SHAP) (version 0.47.1) values were utilized to visualize the contribution of individual lectins and their associated glycan epitopes to OA classification, analyzing the significance of each feature in the model’s predictions and illustrating its impact on the final CatBoost model.

#### Statistical Rationale

Data analysis was performed in R and Excel. Hierarchically clustered heatmaps and volcano plots were generated by the Complexheatmap and ggplot2 packages respectively. Volcano plots were made using the log2 fold change of OA joints relative to control joints for each individual species. GBPs were considered differentially expressed for lectins with a-log10 p-value < 0.05. Linear regression using the LME R package was used to model the association between lectin signal intensity and joint condition (OA vs. control), with sequential models adjusting for age, sex, and age-by-condition interaction. Analyses were performed separately for each species, and lectins showing significant condition effects (p < 0.05).

## Supplementary Figures

**Figure S1:** Heatmap of lectin microarray glycopatterns in synovial fluid from control and osteoarthritic (OA) joints in A) humans (knee), B) horses (carpal (ACJ and MCJ), fetlock, tarsal and stifle), and C) dogs (stifle)

**Figure S2:** Scatterplot of age versus OA grade in equine samples

**Figure S3:** Model performance evaluation for CatBoost classifier Supplementary Materials:

**Table S1-3:** Human, Equine, and Canine demographic data

**Table S4:** Lectins and antibodies highlighted in volcano plots

**Table S5:** Detailed regression results for lectins associated with OA in canine, equine, and human synovial fluid samples

**Table S6:** Pycaret model performances

**Table S7:** Lectin epitope data

**Table S8:** Lectin Microarray workflow

Data S1: Lectin microarray data

This dataset contains normalized fluorescence intensities from the lectin microarray analysis of synovial fluid samples collected from dogs, horses, and humans. Columns include sample ID, condition (Healthy or OA), species, and signal intensities for each lectin probe (e.g., SNA, AMA, BPL, MAA, etc.), along with antibody-based probes (e.g., anti-Lewis, anti-Galectin).

Data S2: Regression outputs for lectin associations with osteoarthritis, age, and sex across canine, equine, and human samples.

This dataset is the full set of regression analyses performed on lectin microarray data, including results with *p* values > 0.05. Columns include regression term (e.g., condition, age, sex), coefficient estimate, standard error, test statistic, *p* value, 95% confidence interval, model specification (unadjusted, adjusted for age, adjusted for age + sex), lectin name, species, and model fit statistics (R², adjusted R², F-statistic, model *p* value).

## Author contributions

Angelo G Peralta: Data curation, Formal analysis, Software, Visualization, Writing – original draft, Writing – review & editing.

Parisa Raeisimakiani: Investigation, Data curation, Visualization. Kei Hayashi: Investigation, Resources, Writing – review & editing.

Lara K Mahal: Resources, Supervision, Visualization, Writing – review & editing. Heidi L Reesink: Conceptualization, Funding acquisition, Project administration, Resources, Supervision, Validation, Writing – review & editing.

## Conflicts of Interest

The authors declare that they have no conflicts of interests with the contents of this article.

## Abbreviations

ACL: anterior cruciate ligament
AIA: *Artocarpus integrifolia* agglutinin
ASF: arthritic synovial fibroblast
AUC: area under the receiver operating characteristic curve
BanLec/H84T: Banana Lectin H84T
CCL: cranial cruciate ligament
CD44: cluster of differentiation 44
CUHA: Cornell University Hospital for Animals
diCBM40: dimeric carbohydrate-binding module 40
GBPs: glycan-binding proteins
HSS: Hospital for Special Surgery
LcH: *Lens culinaris*
IL-6: interleukin-6
IQR: interquartile range
IRB: Institutional Review Board
MCC: Matthews correlation coefficient
MMP: matrix metalloproteinase
MRI: magnetic resonance imaging
OA: osteoarthritis
PHA-E: *Phaseolus vulgaris* lectin E
PTOA: post-traumatic osteoarthritis
ROC: receiver operating characteristic
SHAP: Shapley additive explanations
SLBR-H: Siglec-like binding region *Streptococcus gordonii* strains *DL1*
SLBR-N: Siglec-like binding region *Streptococcus gordonii* strains *UB10712*
SMOTE: Synthetic Minority Oversampling Technique
SNA: *Sambucus nigra* agglutinin
SVM: support vector machine
TGF-β: transforming growth factor beta
TKR: total knee replacement
TLR: Toll-like receptor

## Acknowledgments

The authors thank Camilla Carballo for her contributions to the study. We also thank Scott Rodeo for his contributions to the study and for providing valuable feedback on the manuscript.

This work was supported by the Weill Cornell Medical College/Center for Translational Science NIH Pilot Award (UL1 TR002384), the Harry M. Zweig Memorial Fund for Equine Research, and the Cornell Veterinary Biobank (NIH R24 GM082910).

## Data availability

Supplementary data and code used in this study are available from the corresponding author upon reasonable request

## Supplemental Data

This article contains supplemental data

## Notes

### Competing Interest Statement

The authors have declared no competing interest.

## References

1. Nguyen, A., Lee, P., Rodriguez, E. K., Chahal, K., Freedman, B. R., and Nazarian, A. (2025) Addressing the growing burden of musculoskeletal diseases in the ageing US population: challenges and innovations. Lancet Healthy Longev 6, 100707

2. Maia, C. R., Annichino, R. F., de Azevedo E Souza Munhoz, M., Machado, E. G., Marchi, E., and Castano-Betancourt, M. C. (2023) Post-traumatic osteoarthritis: the worst associated injuries and differences in patients’ profile when compared with primary osteoarthritis. BMC Musculoskelet Disord 24, 568

3. Lawrence, R. C., Felson, D. T., Helmick, C. G., Arnold, L. M., Choi, H., Deyo, R. A., Gabriel, S., Hirsch, R., Hochberg, M. C., Hunder, G. G., Jordan, J. M., Katz, J. N., Kremers, H. M., Wolfe, F., and National Arthritis Data Workgroup (2008) Estimates of the prevalence of arthritis and other rheumatic conditions in the United States. Part II. Arthritis Rheum 58, 26–35

4. Anderson, K. L., O’Neill, D. G., Brodbelt, D. C., Church, D. B., Meeson, R. L., Sargan, D., Summers, J. F., Zulch, H., and Collins, L. M. (2018) Prevalence, duration and risk factors for appendicular osteoarthritis in a UK dog population under primary veterinary care. Sci Rep 8, 5641

5. Baccarin, R. Y. A., Seidel, S. R. T., Michelacci, Y. M., Tokawa, P. K. A., and Oliveira, T. M. (2022) Osteoarthritis: a common disease that should be avoided in the athletic horse’s life. Anim Front 12, 25–36

6. Kosinska, M. K., Mastbergen, S. C., Liebisch, G., Wilhelm, J., Dettmeyer, R. B., Ishaque, B., Rickert, M., Schmitz, G., Lafeber, F. P., and Steinmeyer, J. (2016) Comparative lipidomic analysis of synovial fluid in human and canine osteoarthritis. Osteoarthritis Cartilage 24, 1470–1478

7. Pelletier, J. P., Martel-Pelletier, J., and Abramson, S. B. (2001) Osteoarthritis, an inflammatory disease: potential implication for the selection of new therapeutic targets. Arthritis Rheum 44, 1237–1247

8. Han, M., Dai, J., Zhang, Y., Lin, Q., Jiang, M., Xu, X., and Liu, Q. (2012) Identification of Osteoarthritis Biomarkers by Proteomic Analysis of Synovial Fluid. Journal of International Medical Research,

9. McIlwraith, C. W., Kawcak, C. E., Frisbie, D. D., Little, C. B., Clegg, P. D., Peffers, M. J., Karsdal, M. A., Ekman, S., Laverty, S., Slayden, R. A., Sandell, L. J., Lohmander, L. S., and Kraus, V. B. (2018) Biomarkers for equine joint injury and osteoarthritis. J Orthop Res 36, 823–831

10. Vincent, T. L. (2022) OA synovial fluid: biological insights into a whole-joint disease. Osteoarthritis Cartilage 30, 765–766

11. Carlson, A. K., Rawle, R. A., Wallace, C. W., Brooks, E. G., Adams, E., Greenwood, M. C., Olmer, M., Lotz, M. K., Bothner, B., and June, R. K. (2019) Characterization of synovial fluid metabolomic phenotypes of cartilage morphological changes associated with osteoarthritis. Osteoarthritis Cartilage 27, 1174–1184

12. Mickiewicz, B., Kelly, J. J., Ludwig, T. E., Weljie, A. M., Preston Wiley, J., Schmidt, T. A., and Vogel, H. J. (2015) Metabolic analysis of knee synovial fluid as a potential diagnostic approach for osteoarthritis. Journal of Orthopaedic Research® 33, 1631–1638

13. Noordwijk, K. J., Qin, R., Diaz-Rubio, M. E., Zhang, S., Su, J., Mahal, L. K., and Reesink, H. L. (2022) Metabolism and global protein glycosylation are differentially expressed in healthy and osteoarthritic equine carpal synovial fluid. Equine Vet J 54, 323–333

14. Anderson, J. R., Phelan, M. M., Caamaño-Gutiérrez, E., Clegg, P. D., Rubio-Martinez, L. M., and Peffers, M. J. (2025) Metabolomic and proteomic stratification of equine osteoarthritis. Equine Vet J 57, 1204–1218

15. Altindag, O., Erel, O., Aksoy, N., Selek, S., Celik, H., and Karaoglanoglu, M. (2007) Increased oxidative stress and its relation with collagen metabolism in knee osteoarthritis. Rheumatol Int 27, 339–344

16. Anderson, J. R., Chokesuwattanaskul, S., Phelan, M. M., Welting, T. J. M., Lian, L.-Y., Peffers, M. J., and Wright, H. L. (2018) H NMR Metabolomics Identifies Underlying Inflammatory Pathology in Osteoarthritis and Rheumatoid Arthritis Synovial Joints. J Proteome Res 17, 3780–3790

17. Varki, A. (2017) Biological roles of glycans. Glycobiology 27, 3–49

18. Reily, C., Stewart, T. J., Renfrow, M. B., and Novak, J. (2019) Glycosylation in health and disease. Nat Rev Nephrol 15, 346–366

19. Novokmet, M., Lukić, E., Vučković, F., Ðurić, Ž., Keser, T., Rajšl, K., Remondini, D., Castellani, G., Gašparović, H., Gornik, O., and Lauc, G. (2014) Changes in IgG and total plasma protein glycomes in acute systemic inflammation. Sci Rep 4, 4347

20. Mohideen, F. I., and Mahal, L. K. (2024) Infection and the Glycome─New Insights into Host Response. ACS Infect Dis 10, 2540–2550

21. Turnbull, J. E., and Field, R. A. (2007) Emerging glycomics technologies. Nat Chem Biol 3, 74–77

22. Chen, S., Qin, R., and Mahal, L. K. (2021) Sweet systems: technologies for glycomic analysis and their integration into systems biology. Crit Rev Biochem Mol Biol 56, 301–320

23. Trbojević-Akmačić, I., Lageveen-Kammeijer, G. S. M., Heijs, B., Petrović, T., Deriš, H., Wuhrer, M., and Lauc, G. (2022) High-Throughput Glycomic Methods. Chem Rev 122, 15865–15913

24. Tiemeyer, M., Aoki, K., Paulson, J., Cummings, R. D., York, W. S., Karlsson, N. G., Lisacek, F., Packer, N. H., Campbell, M. P., Aoki, N. P., Fujita, A., Matsubara, M., Shinmachi, D., Tsuchiya, S., Yamada, I., Pierce, M., Ranzinger, R., Narimatsu, H., and Aoki-Kinoshita, K. F. (2017) GlyTouCan: an accessible glycan structure repository. Glycobiology 27, 915–919

25. Smith, D. F., Song, X., and Cummings, R. D. (2010) Use of glycan microarrays to explore specificity of glycan-binding proteins. Methods Enzymol 480, 417–444

26. Pilobello, K. T., Krishnamoorthy, L., Slawek, D., and Mahal, L. K. (2005) Development of a lectin microarray for the rapid analysis of protein glycopatterns. Chembiochem 6, 985–989

27. Bojar, D., and Lisacek, F. (2022) Glycoinformatics in the Artificial Intelligence Era. Chem Rev 122, 15971–15988

28. Sevim Bayrak, C., Forst, C. V., Jones, D. R., Gresham, D. J., Pushalkar, S., Wu, S., Vogel, C., Mahal, L. K., Ghedin, E., Ross, T., García-Sastre, A., and Zhang, B. (2024) Patient subtyping analysis of baseline multi-omic data reveals distinct pre-immune states associated with antibody response to seasonal influenza vaccination. Clin Immunol 266, 110333

29. Ng, S., Masarone, S., Watson, D., and Barnes, M. R. (2023) The benefits and pitfalls of machine learning for biomarker discovery. Cell Tissue Res 394, 17–31

30. Mariethoz, J., Khatib, K., Alocci, D., Campbell, M. P., Karlsson, N. G., Packer, N. H., Mullen, E. H., and Lisacek, F. (2016) SugarBindDB, a resource of glycan-mediated host-pathogen interactions. Nucleic Acids Res 44, D1243–50

31. Deng, X., Liu, X., Zhang, Y., Ke, D., Yan, R., Wang, Q., Tian, X., Li, M., Zeng, X., and Hu, C. (2023) Changes of serum IgG glycosylation patterns in rheumatoid arthritis. Clin Proteomics 20, 7

32. Kissel, T., Toes, R. E. M., Huizinga, T. W. J., and Wuhrer, M. (2023) Glycobiology of rheumatic diseases. Nat Rev Rheumatol 19, 28–43

33. Xu, X., Balmer, L., Chen, Z., Mahara, G., and Lin, L. (2022) The role of IgG N-galactosylation in spondyloarthritis. Transl. Metab. Syndr. Res. 5, 16–23

34. Bhattacharjee, M., Sharma, R., Goel, R., Balakrishnan, L., Renuse, S., Advani, J., Gupta, S. T., Verma, R., Pinto, S. M., Sekhar, N. R., Nair, B., Prasad, T. S. K., Harsha, H. C., Jois, R., Shankar, S., and Pandey, A. (2013) A multilectin affinity approach for comparative glycoprotein profiling of rheumatoid arthritis and spondyloarthropathy. Clin Proteomics 10, 11

35. Kraus, V. B., and Hsueh, M.-F. (2024) Molecular biomarker approaches to prevention of post-traumatic osteoarthritis. Nat Rev Rheumatol 20, 272–289

36. O’Sullivan, O., Ladlow, P., Steiner, K., Hillman, C., Stocks, J., Bennett, A. N., Valdes, A. M., and Kluzek, S. (2023) Current status of catabolic, anabolic and inflammatory biomarkers associated with structural and symptomatic changes in the chronic phase of post-traumatic knee osteoarthritis-a systematic review. Osteoarthr Cartil Open 5, 100412

37. Takase, K., McCulloch, P. C., Yik, J. H. N., and Haudenschild, D. R. (2025) Clinical and molecular landscape of post-traumatic osteoarthritis. Connect Tissue Res, 1–7

38. Yu, H., Li, M., Shu, J., Dang, L., Wu, X., Wang, Y., Wang, X., Chang, X., Bao, X., Zhu, B., Ren, X., Chen, W., and Li, Y. (2023) Characterization of aberrant glycosylation associated with osteoarthritis based on integrated glycomics methods. Arthritis Res Ther 25, 102

39. Liu, H.-Z., Song, X.-Q., and Zhang, H. (2024) Sugar-coated bullets: Unveiling the enigmatic mystery “sweet arsenal” in osteoarthritis. Heliyon 10, e27624

40. Fuehrer, J., Pichler, K. M., Fischer, A., Giurea, A., Weinmann, D., Altmann, F., Windhager, R., Gabius, H.-J., and Toegel, S. (2021) N-Glycan profiling of chondrocytes and fibroblast-like synoviocytes: Towards functional glycomics in osteoarthritis. Proteomics Clin Appl 15, e2000057

41. Luo, Y., Wu, Z., Chen, S., Luo, H., Mo, X., Wang, Y., and Tang, J. (2022) Protein N-glycosylation aberrations and glycoproteomic network alterations in osteoarthritis and osteoarthritis with type 2 diabetes. Sci Rep 12, 6977

42. Heindel, D. W., Koppolu, S., Zhang, Y., Kasper, B., Meche, L., Vaiana, C. A., Bissel, S. J., Carter, C. E., Kelvin, A. A., Elaish, M., Lopez-Orozco, J., Zhang, B., Zhou, B., Chou, T.-W., Lashua, L., Hobman, T. C., Ross, T. M., Ghedin, E., and Mahal, L. K. (2020) Glycomic analysis of host response reveals high mannose as a key mediator of influenza severity. Proc Natl Acad Sci U S A 117, 26926–26935

43. Heindel, D. W., Chen, S., Aziz, P. V., Chung, J. Y., Marth, J. D., and Mahal, L. K. (2022) Glycomic Analysis Reveals a Conserved Response to Bacterial Sepsis Induced by Different Bacterial Pathogens. ACS Infect Dis 8, 1075–1085

44. Kurz, E., Chen, S., Vucic, E., Baptiste, G., Loomis, C., Agrawal, P., Hajdu, C., Bar-Sagi, D., and Mahal, L. K. (2021) Integrated Systems Analysis of the Murine and Human Pancreatic Cancer Glycomes Reveals a Tumor-Promoting Role for ST6GAL1. Mol Cell Proteomics 20, 100160

45. Pilobello, K. T., Slawek, D. E., and Mahal, L. K. (2007) A ratiometric lectin microarray approach to analysis of the dynamic mammalian glycome. Proceedings of the National Academy of Sciences 104, 11534–11539

46. York, W. S., Agravat, S., Aoki-Kinoshita, K. F., McBride, R., Campbell, M. P., Costello, C. E., Dell, A., Feizi, T., Haslam, S. M., Karlsson, N., Khoo, K.-H., Kolarich, D., Liu, Y., Novotny, M., Packer, N. H., Paulson, J. C., Rapp, E., Ranzinger, R., Rudd, P. M., Smith, D. F., Struwe, W. B., Tiemeyer, M., Wells, L., Zaia, J., and Kettner, C. (2014) MIRAGE: the minimum information required for a glycomics experiment. Glycobiology 24, 402–406

47. Tateno, H., Mahal, L. K., Feizi, T., Kettner, C., and Paulson, J. C. (2025) The minimum information required for a glycomics experiment (MIRAGE) project: improving the standards for reporting lectin microarray data. Glycobiology 35,

48. Srivastava, S., Verhagen, A., Sasmal, A., Wasik, B. R., Diaz, S., Yu, H., Bensing, B. A., Khan, N., Khedri, Z., Secrest, P., Sullam, P., Varki, N., Chen, X., Parrish, C. R., and Varki, A. (2022) Development and applications of sialoglycan-recognizing probes (SGRPs) with defined specificities: exploring the dynamic mammalian sialoglycome. Glycobiology 32, 1116–1136

49. Bensing, B. A., Li, Q., Park, D., Lebrilla, C. B., and Sullam, P. M. (2018) Streptococcal Siglec-like adhesins recognize different subsets of human plasma glycoproteins: implications for infective endocarditis. Glycobiology 28, 601–611

50. Bensing, B. A., Khedri, Z., Deng, L., Yu, H., Prakobphol, A., Fisher, S. J., Chen, X., Iverson, T. M., Varki, A., and Sullam, P. M. (2016) Novel aspects of sialoglycan recognition by the Siglec-like domains of streptococcal SRR glycoproteins. Glycobiology 26, 1222–1234

51. Zhu, W., Zhou, Y., Guo, L., and Feng, S. (2024) Biological function of sialic acid and sialylation in human health and disease. Cell Death Discov 10, 415

52. Li, Y., and Chen, X. (2012) Sialic acid metabolism and sialyltransferases: natural functions and applications. Appl Microbiol Biotechnol 94, 887–905

53. Varki, A. (2008) Sialic acids in human health and disease. Trends Mol Med 14, 351–360

54. Lee, H., Lee, A., Seo, N., Oh, J., Kweon, O.-K., An, H. J., and Kim, J. (2020) Discovery of N-glycan Biomarkers for the Canine Osteoarthritis. Life (Basel) 10,

55. Svala, E., Jin, C., Rüetschi, U., Ekman, S., Lindahl, A., Karlsson, N. G., and Skiöldebrand, E. (2017) Characterisation of lubricin in synovial fluid from horses with osteoarthritis. Equine Vet J 49, 116–123

56. Shibuya, N., Goldstein, I. J., Broekaert, W. F., Nsimba-Lubaki, M., Peeters, B., and Peumans, W. J. (1987) The elderberry (Sambucus nigra L.) bark lectin recognizes the Neu5Ac(alpha 2-6)Gal/GalNAc sequence. J Biol Chem 262, 1596–1601

57. Geisler, C., and Jarvis, D. L. (2011) Effective glycoanalysis with Maackia amurensis lectins requires a clear understanding of their binding specificities. Glycobiology 21, 988–993

58. Qin, R., Kurz, E., Chen, S., Zeck, B., Chiribogas, L., Jackson, D., Herchen, A., Attia, T., Carlock, M., Rapkiewicz, A., Bar-Sagi, D., Ritchie, B., Ross, T. M., and Mahal, L. K. (2022) α2,6-Sialylation Is Upregulated in Severe COVID-19, Implicating the Complement Cascade. ACS Infect Dis 8, 2348–2361

59. Wang, X., Inoue, S., Gu, J., Miyoshi, E., Noda, K., Li, W., Mizuno-Horikawa, Y., Nakano, M., Asahi, M., Takahashi, M., Uozumi, N., Ihara, S., Lee, S. H., Ikeda, Y., Yamaguchi, Y., Aze, Y., Tomiyama, Y., Fujii, J., Suzuki, K., Kondo, A., Shapiro, S. D., Lopez-Otin, C., Kuwaki, T., Okabe, M., Honke, K., and Taniguchi, N. (2005) Dysregulation of TGF-beta1 receptor activation leads to abnormal lung development and emphysema-like phenotype in core fucose-deficient mice. Proc Natl Acad Sci U S A 102, 15791–15796

60. Wilson, J. R., Williams, D., and Schachter, H. (1976) The control of glycoprotein synthesis: N-acetylglucosamine linkage to a mannose residue as a signal for the attachment of L-fucose to the asparagine-linked N-acetylglucosamine residue of glycopeptide from alpha1-acid glycoprotein. Biochem Biophys Res Commun 72, 909–916

61. Wang, Y., Yuan, R., Liang, B., Zhang, J., Wen, Q., Chen, H., Tian, Y., Wen, L., and Zhou, H. (2024) A “One-Step” Strategy for the Global Characterization of Core-Fucosylated Glycoproteome. JACS Au 4, 2005–2018

62. Zhang, N.-Z., Zhao, L.-F., Zhang, Q., Fang, H., Song, W.-L., Li, W.-Z., Ge, Y.-S., and Gao, P. (2023) Core fucosylation and its roles in gastrointestinal glycoimmunology. World J Gastrointest Oncol 15, 1119–1134

63. García-García, A., Serna, S., Yang, Z., Delso, I., Taleb, V., Hicks, T., Artschwager, R., Vakhrushev, S. Y., Clausen, H., Angulo, J., Corzana, F., Reichardt, N. C., and Hurtado-Guerrero, R. (2021) FUT8-Directed Core Fucosylation of N-glycans Is Regulated by the Glycan Structure and Protein Environment. ACS Catal 11, 9052– 9065

64. Howard, I. K., Sage, H. J., Stein, M. D., Young, N. M., Leon, M. A., and Dyckes, D. F. (1971) Studies on a phytohemagglutinin from the lentil. II. Multiple forms of Lens culinaris hemagglutinin. J Biol Chem 246, 1590–1595

65. Kochibe, N., and Furukawa, K. (1980) Purification and properties of a novel fucose-specific hemagglutinin of Aleuria aurantia. Biochemistry 19, 2841–2846

66. Brockhausen, I., Schachter, H., and Stanley, P. (2009) in Essentials of Glycobiology (Cold Spring Harbor Laboratory Press, Cold Spring Harbor (NY)).

67. Ali, L., Flowers, S. A., Jin, C., Bennet, E. P., Ekwall, A.-K. H., and Karlsson, N. G. (2014) The O-glycomap of lubricin, a novel mucin responsible for joint lubrication, identified by site-specific glycopeptide analysis. Mol Cell Proteomics 13, 3396–3409

68. Boushehri, S., Holey, H., Brosz, M., Gumbsch, P., Pastewka, L., Aponte-Santamaría, C., and Gräter, F. (2024) O-glycans Expand Lubricin and Attenuate Its Viscosity and Shear Thinning. Biomacromolecules 25, 3893–3908

69. Afshari, A. R., Chang, V., Thomsson, K. A., Höglund, J., Browne, E. N., Karadzhov, G., Mahoney, K. E., Lucas, T. M., Rangel-Angarita, V., Ryberg, H., Gidwani, K., Pettersson, K., Rolfson, O., Björkman, L. I., Eisler, T., Schmidt, T. A., Jay, G. D., Malaker, S. A., and Karlsson, N. G. (2025) Glycoproteoforms of Osteoarthritis-associated Lubricin in Plasma and Synovial Fluid. Mol Cell Proteomics 24, 100923

70. Jay, G. D., Harris, D. A., and Cha, C. J. (2001) Boundary lubrication by lubricin is mediated by O-linked beta(1-3)Gal-GalNAc oligosaccharides. Glycoconj J 18, 807– 815

71. Wang, Y., Gludish, D. W., Hayashi, K., Todhunter, R. J., Krotscheck, U., Johnson, P. J., Cummings, B. P., Su, J., and Reesink, H. L. (2020) Synovial fluid lubricin increases in spontaneous canine cruciate ligament rupture. Sci Rep 10, 16725

72. Bausch, J. N., and Poretz, R. D. (1977) Purification and properties of the hemagglutinin from Maclura pomifera seeds. Biochemistry 16, 5790–5794

73. Kabir, S., and Daar, A. S. (1994) The composition and properties of jacalin, a lectin of diverse applications obtained from the jackfruit (Artocarpus heterophyllus) seeds. Immunol Invest 23, 167–188

74. Bojar, D., Meche, L., Meng, G., Eng, W., Smith, D. F., Cummings, R. D., and Mahal, L. K. (2022) A Useful Guide to Lectin Binding: Machine-Learning Directed Annotation of 57 Unique Lectin Specificities. ACS Chem Biol 17, 2993–3012

75. Watkins, A. R., and Reesink, H. L. (2020) Lubricin in experimental and naturally occurring osteoarthritis: a systematic review. Osteoarthritis Cartilage 28, 1303–1315

76. Neu, C. P., Reddi, A. H., Komvopoulos, K., Schmid, T. M., and Di Cesare, P. E. (2010) Increased friction coefficient and superficial zone protein expression in patients with advanced osteoarthritis. Arthritis Rheum 62, 2680–2687

77. Maki, Y., Otani, Y., Okamoto, R., Izumi, M., and Kajihara, Y. (2022) Isolation and characterization of high-mannose type glycans containing five or six mannose residues from hen egg yolk. Carbohydr Res 521, 108680

78. Šèupáková, K., Adelaja, O. T., Balluff, B., Ayyappan, V., Tressler, C. M., Jenkinson, N. M., Claes, B., Sr, Bowman, A. P., Cimino-Mathews, A. M., White, M. J., Argani, P., Heeren, R. M., and Glunde, K. (2021) Clinical importance of high-mannose, fucosylated, and complex N-glycans in breast cancer metastasis. JCI Insight 6,

79. Boyaval, F., Dalebout, H., Van Zeijl, R., Wang, W., Fariña-Sarasqueta, A., Lageveen-Kammeijer, G. S. M., Boonstra, J. J., McDonnell, L. A., Wuhrer, M., Morreau, H., and Heijs, B. (2022) High-Mannose-Glycans as Malignant Progression Markers in Early-Stage Colorectal Cancer. Cancers (Basel) 14,

80. Tsui, C. K., Twells, N., Durieux, J., Doan, E., Woo, J., Khosrojerdi, N., Brooks, J., Kulepa, A., Webster, B., Mahal, L. K., and Dillin, A. (2024) CRISPR screens and lectin microarrays identify high mannose N-glycan regulators. Nat Commun 15, 9970

81. Lusvarghi, S., and Bewley, C. A. (2016) Griffithsin: An Antiviral Lectin with Outstanding Therapeutic Potential. Viruses 8,

82. Koshte, V. L., van Dijk, W., van der Stelt, M. E., and Aalberse, R. C. (1990) Isolation and characterization of BanLec-I, a mannoside-binding lectin from Musa paradisiac (banana). Biochem J 272, 721–726

83. Covés-Datson, E. M., King, S. R., Legendre, M., Gupta, A., Chan, S. M., Gitlin, E., Kulkarni, V. V., Pantaleón García, J., Smee, D. F., Lipka, E., Evans, S. E., Tarbet, E. B., Ono, A., and Markovitz, D. M. (2020) A molecularly engineered antiviral banana lectin inhibits fusion and is efficacious against influenza virus infection in vivo. Proc Natl Acad Sci U S A 117, 2122–2132

84. Metcalfe, A. J., Andersson, M. L. E., Goodfellow, R., and Thorstensson, C. A. (2012) Is knee osteoarthritis a symmetrical disease? Analysis of a 12 year prospective cohort study. BMC Musculoskelet Disord 13, 153

85. Vanhooren, V., Dewaele, S., Libert, C., Engelborghs, S., De Deyn, P. P., Toussaint, O., Debacq-Chainiaux, F., Poulain, M., Glupczynski, Y., Franceschi, C., Jaspers, K., van der Pluijm, I., Hoeijmakers, J., and Chen, C. C. (2010) Serum N-glycan profile shift during human ageing. Exp Gerontol 45, 738–743

86. Ding, N., Nie, H., Sun, X., Sun, W., Qu, Y., Liu, X., Yao, Y., Liang, X., Chen, C. C., and Li, Y. (2011) Human serum N-glycan profiles are age and sex dependent. Age Ageing 40, 568–575

87. Li, H., Patel, V., DiMartino, S. E., Froehlich, J. W., and Lee, R. S. (2020) An in-depth Comparison of the Pediatric and Adult Urinary N-glycomes. Mol Cell Proteomics 19, 1767–1776

88. Ali, M. (2020) PyCaret: An open-source, low-code machine learning library in Python.

89. Cai, Y., Yuan, Y., and Zhou, A. (2024) Predictive slope stability early warning model based on CatBoost. Sci Rep 14, 25727

90. Xin, Z.-C., Zhang, J.-S., and Liu, Q. (2025) Predicting CaO activity in multiple slag system using improved whale optimization algorithm and categorical boosting. Sci Rep 15, 9533

91. Lundberg, S. M., Erion, G., Chen, H., DeGrave, A., Prutkin, J. M., Nair, B., Katz, R., Himmelfarb, J., Bansal, N., and Lee, S.-I. (2020) From Local Explanations to Global Understanding with Explainable AI for Trees. Nat Mach Intell 2, 56–67

92. Lundberg, S., and Lee, S.-I. (2017) A unified approach to interpreting model predictions. arXiv [cs.AI],

93. Wang, Y., Pan, P., Khan, A., Çil, Ç., and Pineda, M. A. (2022) Synovial Fibroblast Sialylation Regulates Cell Migration and Activation of Inflammatory Pathways in Arthritogenesis. Front Immunol 13, 847581

94. Flowers, S. A., Thomsson, K. A., Ali, L., Huang, S., Mthembu, Y., Regmi, S. C., Holgersson, J., Schmidt, T. A., Rolfson, O., Björkman, L. I., Sundqvist, M., Karlsson-Bengtsson, A., Jay, G. D., Eisler, T., Krawetz, R., and Karlsson, N. G. (2020) Decrease of core 2 glycans on synovial lubricin in osteoarthritis reduces galectin-3 mediated crosslinking. J Biol Chem 295, 16023–16036

95. Silva, A. D., Hwang, J., Marciel, M. P., and Bellis, S. L. (2024) The pro-inflammatory cytokines IL-1β and IL-6 promote upregulation of the ST6GAL1 sialyltransferase in pancreatic cancer cells. J Biol Chem 300, 107752

96. Varki, A., Freeze, H. H., and Gagneux, P. (2009) in Essentials of Glycobiology, eds Varki A, Cummings RD, Esko JD, Freeze HH, Stanley P, Bertozzi CR, Hart GW, Etzler ME (Cold Spring Harbor Laboratory Press, Cold Spring Harbor (NY)).

97. Al-Sharif, A., Jamal, M., Zhang, L. X., Larson, K., Schmidt, T. A., Jay, G. D., and Elsaid, K. A. (2015) Lubricin/Proteoglycan 4 Binding to CD44 Receptor: A Mechanism of the Suppression of Proinflammatory Cytokine-Induced Synoviocyte Proliferation by Lubricin. Arthritis Rheumatol 67, 1503–1513

98. Iqbal, S. M., Leonard, C., Regmi, S. C., De Rantere, D., Tailor, P., Ren, G., Ishida, H., Hsu, C., Abubacker, S., Pang, D. S., Salo, P. T., Vogel, H. J., Hart, D. A., Waterhouse, C. C., Jay, G. D., Schmidt, T. A., and Krawetz, R. J. (2016) Lubricin/Proteoglycan 4 binds to and regulates the activity of Toll-Like Receptors In Vitro. Sci Rep 6, 18910

99. Homan, K., Onodera, T., Hanamatsu, H., Furukawa, J.-I., Momma, D., Matsuoka, M., and Iwasaki, N. (2023) Articular cartilage corefucosylation regulates tissue resilience in osteoarthritis. eLife 12,

100. Gao, C., Maeno, T., Ota, F., Ueno, M., Korekane, H., Takamatsu, S., Shirato, K., Matsumoto, A., Kobayashi, S., Yoshida, K., Kitazume, S., Ohtsubo, K., Betsuyaku, T., and Taniguchi, N. (2012) Sensitivity of heterozygous α1,6-fucosyltransferase knock-out mice to cigarette smoke-induced emphysema: implication of aberrant transforming growth factor-β signaling and matrix metalloproteinase gene expression. J Biol Chem 287, 16699–16708

101. Urita, A., Matsuhashi, T., Onodera, T., Nakagawa, H., Hato, M., Amano, M., Seito, N., Minami, A., Nishimura, S.-I., and Iwasaki, N. (2011) Alterations of high-mannose type N-glycosylation in human and mouse osteoarthritis cartilage. Arthritis Rheum 63, 3428–3438

102. Lopandić, Z., Dragačević, L., Popović, D., Andjelković, U., Minić, R., and Gavrović-Jankulović, M. (2021) BanLec-eGFP Chimera as a Tool for Evaluation of Lectin Binding to High-Mannose Glycans on Microorganisms. Biomolecules 11,

103. Mori, T., O’Keefe, B. R., Sowder, R. C., 2nd, Bringans, S., Gardella, R., Berg, S., Cochran, P., Turpin, J. A., Buckheit, R. W., Jr, McMahon, J. B., and Boyd, M. R. (2005) Isolation and characterization of griffithsin, a novel HIV-inactivating protein, from the red alga Griffithsia sp. J Biol Chem 280, 9345–9353

104. Zhang, N., Lin, S., Cui, W., and Newman, P. J. (2022) Overlapping and unique substrate specificities of ST3GAL1 and 2 during hematopoietic and megakaryocytic differentiation. Blood Adv 6, 3945–3955

105. Comelli, E. M., Head, S. R., Gilmartin, T., Whisenant, T., Haslam, S. M., North, S. J., Wong, N.-K., Kudo, T., Narimatsu, H., Esko, J. D., Drickamer, K., Dell, A., and Paulson, J. C. (2006) A focused microarray approach to functional glycomics: transcriptional regulation of the glycome. Glycobiology 16, 117–131

106. Krištić, J., Vučković, F., Menni, C., Klarić, L., Keser, T., Beceheli, I., Pučić-Baković, M., Novokmet, M., Mangino, M., Thaqi, K., Rudan, P., Novokmet, N., Sarac, J., Missoni, S., Kolčić, I., Polašek, O., Rudan, I., Campbell, H., Hayward, C., Aulchenko, Y., Valdes, A., Wilson, J. F., Gornik, O., Primorac, D., Zoldoš, V., Spector, T., and Lauc, G. (2014) Glycans are a novel biomarker of chronological and biological ages. J Gerontol A Biol Sci Med Sci 69, 779–789

107. Han, J., Pan, Y., Qin, W., Gu, Y., Xu, X., Zhao, R., Sha, J., Zhang, R., Gu, J., and Ren, S. (2020) Quantitation of sex-specific serum N-glycome changes in expression level during mouse aging based on Bionic Glycome method. Exp Gerontol 141, 111098

108. Sanchez-Martinez, S., Camara, O., Piella, G., Cikes, M., González-Ballester, M. Á., Miron, M., Vellido, A., Gómez, E., Fraser, A. G., and Bijnens, B. (2021) Machine Learning for Clinical Decision-Making: Challenges and Opportunities in Cardiovascular Imaging. Front Cardiovasc Med 8, 765693

109. Jordan, M. I., and Mitchell, T. M. (2015) Machine learning: Trends, perspectives, and prospects. Science 349, 255–260

110. Rajula, H. S. R., Verlato, G., Manchia, M., Antonucci, N., and Fanos, V. (2020) Comparison of Conventional Statistical Methods with Machine Learning in Medicine: Diagnosis, Drug Development, and Treatment. Medicina (Kaunas) 56,

111. Srinivasu, P. N., Jaya Lakshmi, G., Gudipalli, A., Narahari, S. C., Shafi, J., Woźniak, M., and Ijaz, M. F. (2024) XAI-driven CatBoost multi-layer perceptron neural network for analyzing breast cancer. Scientific Reports 14, 1–19

112. Gonca, M., Gul, B. B., and Sert, M. F. (2024) How successful is the CatBoost classifier in diagnosing different dental anomalies in patients via sella turcica and vertebral morphologic alteration? BMC Med Inform Decis Mak 24, 237

113. Hanani, A. A., Donmez, T. B., Kutlu, M., and Mansour, M. (2025) Predicting thyroid cancer recurrence using supervised CatBoost: A SHAP-based explainable AI approach. Medicine (Baltimore) 104, e42667

114. Hashimoto, K., Tokimatsu, T., Kawano, S., Yoshizawa, A. C., Okuda, S., Goto, S., and Kanehisa, M. (2009) Comprehensive analysis of glycosyltransferases in eukaryotic genomes for structural and functional characterization of glycans. Carbohydr Res 344, 881–887

115. Raju, T. S., Briggs, J. B., Borge, S. M., and Jones, A. J. (2000) Species-specific variation in glycosylation of IgG: evidence for the species-specific sialylation and branch-specific galactosylation and importance for engineering recombinant glycoprotein therapeutics. Glycobiology 10, 477–486

116. Medzihradszky, K. F., Kaasik, K., and Chalkley, R. J. (2015) Tissue-Specific Glycosylation at the Glycopeptide Level. Mol Cell Proteomics 14, 2103–2110

117. Aoki-Kinoshita, K. F. (2016) A Practical Guide to Using Glycomics Databases (Springer)

118. Raman, R., Raguram, S., Venkataraman, G., Paulson, J. C., and Sasisekharan, R. (2005) Glycomics: an integrated systems approach to structure-function relationships of glycans. Nat Methods 2, 817–824

119. Rudd, P. M., Karlsson, N. G., Khoo, K.-H., Thaysen-Andersen, M., Wells, L., and Packer, N. H. (2022) in Essentials of Glycobiology, eds Varki A, Cummings RD, Esko JD, Stanley P, Hart GW, Aebi M, Mohnen D, Kinoshita T, Packer NH, Prestegard JH, Schnaar RL, Seeberger PH (Cold Spring Harbor Laboratory Press, Cold Spring Harbor (NY)).

120. Song, X., Heimburg-Molinaro, J., Smith, D. F., and Cummings, R. D. (2015) Glycan microarrays of fluorescently-tagged natural glycans. Glycoconj J 32, 465– 473

121. Loke, I., Kolarich, D., Packer, N. H., and Thaysen-Andersen, M. (2016) Emerging roles of protein mannosylation in inflammation and infection. Mol Aspects Med 51, 31–55

122. Womack, S. J., Carballo, C. B., Secor, E. J., Rodeo, S. A., and Reesink, H. L. (2025) Proteomics Reveals Increased Periostin in Synovial Fluid From Canine and Human Anterior Cruciate Ligament Injury. J Orthop Res 43, 1239–1249

123. Zeni, J. A., Jr, and Snyder-Mackler, L. (2010) Most patients gain weight in the 2 years after total knee arthroplasty: comparison to a healthy control group. Osteoarthritis Cartilage 18, 510–514

124. Tschon, M., Contartese, D., Pagani, S., Borsari, V., and Fini, M. (2021) Gender and Sex Are Key Determinants in Osteoarthritis Not Only Confounding Variables. A Systematic Review of Clinical Data. J Clin Med 10,

125. Colbath, A., and Haubruck, P. (2023) Closing the gap: sex-related differences in osteoarthritis and the ongoing need for translational studies. Ann Transl Med 11, 339

126. Stewart, H. L., Gilbert, D., Stefanovski, D., Garman, Z., Albro, M. B., Bais, M., Grinstaff, M. W., Snyder, B. D., and Schaer, T. P. (2024) A missed opportunity: A scoping review of the effect of sex and age on osteoarthritis using large animal models. Osteoarthritis Cartilage 32, 501–513

127. Anderson, K. L., Zulch, H., O’Neill, D. G., Meeson, R. L., and Collins, L. M. (2020) Risk Factors for Canine Osteoarthritis and Its Predisposing Arthropathies: A Systematic Review. Front Vet Sci 7, 220

128. Kol, A., Arzi, B., Athanasiou, K. A., Farmer, D. L., Nolta, J. A., Rebhun, R. B., Chen, X., Griffiths, L. G., Verstraete, F. J. M., Murphy, C. J., and Borjesson, D. L. (2015) Companion animals: Translational scientist’s new best friends. Sci Transl Med 7, 308ps21

129. Gargiulo, S., Vecchiarelli, L., Pagni, E., and Gramanzini, M. (2025) The Role of Canine Models of Human Cancer: Overcoming Drug Resistance Through a Transdisciplinary “One Health, One Medicine” Approach. Cancers (Basel) 17,

130. Dolnicka, A., Fosse, V., Raciborska, A., and Śmieszek, A. (2025) Building a Therapeutic Bridge Between Dogs and Humans: A Review of Potential Cross-Species Osteosarcoma Biomarkers. Int J Mol Sci 26,

131. McCue, M. E., and McCoy, A. M. (2017) The Scope of Big Data in One Medicine: Unprecedented Opportunities and Challenges. Front Vet Sci 4, 194

132. Koppolu, S., Wang, L., Mathur, A., Nigam, J. A., Dezzutti, C. S., Isaacs, C., Meyn, L., Bunge, K. E., Moncla, B. J., Hillier, S. L., Rohan, L. C., and Mahal, L. K. (2018) Vaginal Product Formulation Alters the Innate Antiviral Activity and Glycome of Cervicovaginal Fluids with Implications for Viral Susceptibility. ACS Infect Dis 4, 1613–1622

