## Supplementary Data for "Lectin Microarray-based Glycomics and Machine Learning Identify Shared Osteoarthritis Biomarkers in Humans, Dogs, and Horses"

### Supplemental Materials

#### Lectin Microarray-Based Glycosylation Profiling and Explainable Machine Learning Reveal Shared Synovial Fluid Osteoarthritis Biomarkers in Humans and Companion Animals

Angelo G Peralta[1], Parisa Raeisimakiani[2], Kei Hayashi[3], Lara K Mahal[2], Heidi L Reesink[1,3,*]

[1]School of Veterinary Medicine, University of California, Davis, Davis CA 95626 USA

[2]Department of Chemistry, University of Alberta, AB T6G 2N4 Canada

[3]College of Veterinary Medicine, Cornell University, Ithaca NY 14853 USA

*Corresponding

### Table of Contents

**Table S1.** Human cohort demographics

**Table S2.** Equine cohort demographics

**Table S3.** Canine cohort demographics

**Table S4.** Lectins and antibodies highlighted in volcano plots

**Table S5.** Lectin panel and glycan binding specificities

**Table S6.** Machine learning model performance metrics

**Table S7.** Additional experimental information for lectin microarray

**Table S8.** Lectin microarray workflow details

**Figure S1.** Heatmap of lectin microarray glycopatterns

**Figure S2.** Scatterplot of age versus OA grade in equine samples

**Figure S3.** Model performance evaluation for CatBoost classifier

**Data S1.** Lectin microarray processed dataset (Excel file, separate)

**Data S2.** Regression analysis dataset (Excel file, separate)

Table S1: Demographic details of the human cohort.

Human synovial fluid (SF) samples were obtained from patients with post-traumatic osteoarthritis (PTOA) secondary to anterior cruciate ligament (ACL) injury or from patients undergoing total knee replacement (TKR) for idiopathic osteoarthritis (OA). Values include sample ID, group, sex, and age. Summary statistics (range, median, mean, standard error [SE], and interquartile range [IQR]) are provided. Abbreviations: ACL, anterior cruciate ligament; TKR, total knee replacement; OA, osteoarthritis

| **Control** | | | | **OA** | | | |
|---|---|---|---|---|---|---|---|
| No. | Group | Sex | Age | No. | Group | Sex | Age |
| HE10 | TKR | F | 63 | OA1 | TKR | M | 63 |
| HE11 | ACL | F | 64 | OA10 | TKR | F | 63 |
| HE12 | TKR | M | 75 | OA11 | ACL | F | 64 |
| HE13 | TKR | M | 78 | OA12 | TKR | M | 75 |
| HE17 | TKR | M | 68 | OA13 | TKR | M | 78 |
| HE18 | ACL | M | 40 | OA14 | ACL | M | 18 |
| HE19 | ACL | F | 18 | OA15 | ACL | F | 45 |
| HE20 | ACL | F | 42 | OA16 | ACL | F | 21 |
| HE21 | TKR | M | 66 | OA17 | TKR | M | 68 |
| HE22 | ACL | F | 26 | OA18 | ACL | M | 40 |
| HE23 | ACL | F | 51 | OA19 | ACL | F | 18 |
| HE24 | ACL | M | 48 | OA2 | TKR | M | 59 |
| HE25 | ACL | M | 28 | OA20 | ACL | F | 42 |
| HE26 | ACL | M | 22 | OA21 | TKR | M | 66 |
| HE8 | TKR | F | 71 | OA22 | ACL | F | 26 |
| HE9 | ACL | M | 29 | OA23 | ACL | F | 51 |
| Range | **38% TKR 62% ACL** | **56% M 44% F** | 18-78 | OA24 | ACL | M | 48 |
| Median | | | 49.5 | OA25 | ACL | M | 28 |
| Mean | | | 49.3 | OA26 | ACL | M | 22 |
| SE | | | 5.1 | OA27 | ACL | M | 24 |
| IQR | | | 36.5 | OA28 | ACL | M | 30 |
| | | | | OA29 | ACL | F | 31 |
| | | | | OA3 | TKR | M | 59 |
| | | | | OA30 | ACL | M | 32 |
| | | | | OA31 | ACL | F | 18 |

| | | | |
|---|---|---|---|
| OA32 | ACL | F | 22 |
| OA33 | ACL | M | 20 |
| OA34 | ACL | M | 38 |
| OA35 | ACL | F | 33 |
| OA36 | ACL | M | 23 |
| OA4 | TKR | M | 64 |
| OA5 | ACL | M | 15 |
| OA6 | ACL | F | 25 |
| OA7 | TKR | M | 62 |
| OA8 | TKR | F | 71 |
| OA9 | ACL | M | 29 |
| Range | **31% TKR 69% ACL** | **61% M 39% F** | 15-78 |
| Median | | | 35.5 |
| Mean | | | 41.4 |
| SE | | | 3.3 |
| IQR | | | 37 |

Table S2. Demographic details of the equine cohort.

Equine synovial fluid samples were obtained from horses with PTOA involving various joints, and healthy controls. Values include sample ID, breed, sex, age, and joint sampled. Summary statistics (range, median, mean, SE, and IQR) are provided. Abbreviations: MCJ, middle carpal joint; ACJ, antebrachiocarpal joint; MCPJ, metacarpophalangeal joint; MTPJ, metatarsophalangeal joint; TCJ, tarsocrural joint; QH, Quarter Horse; TB, Thoroughbred; STB, Standardbred.

| **Control** | | | | | **OA** | | | | |
|---|---|---|---|---|---|---|---|---|---|
| No. | Breed | Sex | Age | Joint | No. | Breed | Sex | Age | Joint |
| 119 | WB | F | 2 | MCJ | 32 | TB | MC | 3 | Carpus |
| 93 | Mixed | MC | 5 | MCJ | 57 | TB | M | 2 | MCJ |
| 11 | TB | MC | 3 | MCJ | 33 | TB | MC | 17 | Carpus |
| 118 | TB | F | 4 | MCPJ | 75 | TB | F | 2 | ACJ |
| 88 | TB | F | 4 | ACJ | 1 | TB | MC | 3 | MCJ |
| 89 | TB | F | 9 | ACJ | 66 | TB | F | 4 | MCJ |
| 9 | TB | F | 4 | MCJ | 41 | STB | MC | 2 | MCJ |
| 16 | TB | F | 4 | MCJ | 55 | TB | F | 2 | MCJ |
| 117 | TB | F | 6 | MCPJ | 31 | Mixed | MC | 6 | Fetlock |
| 10 | TB | F | 3 | MCJ | 58 | TB | F | 4 | MCJ |
| 101 | QH | MC | 3 | ACJ | 63 | TB | F | 3 | MCJ |
| 102 | STB | MC | 3 | MCJ | 65 | TB | F | 3 | ACJ |
| 12 | Paint | M | 2 | MCJ | 45 | STB | M | 3 | MCJ |
| 13 | TB | MC | 5 | MCJ | 19 | TB | MC | 0.7 | MTPJ |
| 100 | STB | MC | 3 | MCJ | 22 | TB | MC | 5 | Carpus |
| 115 | STB | MC | 1 | MCPJ | 8 | TB | MC | 2 | MCJ |
| 96 | Mixed | F | 5 | MCJ | 24 | TB | MC | 12 | Fetlock |
| 106 | Saddlebred | MC | 4 | ACJ | 70 | STB | MC | 3 | MCJ |
| 92 | Mixed | F | 5 | MCJ | 30 | TB | MC | 2 | MCJ |
| 104 | Saddlebred | F | 7 | ACJ | 26 | TB | MC | 11 | Fetlock |
| 108 | TB | F | 4 | TCJ | 25 | STB | MC | 9 | MCPJ |
| 87 | Paint | M | 2 | ACJ | 6 | TB | MC | 6 | MCJ |
| 111 | TB | MC | 15 | TCJ | 77 | QH | MC | 13 | ACJ |
| 114 | STB | MC | 5 | MCPJ | 23 | TB | MC | 2 | |
| 110 | QH | MC | 14 | MCPJ | 21 | TB | F | 3 | ACJ |

| | | | | | | | | | |
|---|---|---|---|---|---|---|---|---|---|
| 90 | TB | MC | 10 | MCJ | 53 | TB | F | 2 | MCJ |
| 103 | TB | F | 5 | MCJ | 27 | TB | F | 4 | Fetlock |
| 94 | Mixed | F | 5 | MCJ | 17 | TB | MC | 3 | ACJ |
| 116 | TB | F | 6 | TCJ | 18 | TB | F | 2 | ACJ |
| 95 | Mixed | F | 5 | MCJ | 67 | TB | MC | 2 | MCJ |
| 109 | QH | F | 4 | MCPJ | 59 | TB | M | 3 | ACJ |
| 98 | TB | F | 9 | ACJ | 40 | TB | MC | 3 | ACJ |
| 97 | TB | F | 4 | ACJ | 34 | TB | MC | 7 | Tarsus |
| 15 | TB | MC | 2 | MCJ | 4 | TB | F | 4 | |
| 91 | QH | F | 7 | ACJ | 71 | QH | F | 13 | ACJ |
| 99 | STB | MC | 5 | ACJ | 50 | TB | F | 2 | MCJ |
| 112 | QH | MC | 12 | Stifle | 81 | TB | MC | 3 | MCJ |
| 14 | TB | MC | 2 | MCJ | 5 | TB | M | 3 | MCJ |
| 113 | STB | M | 1 | MTPJ | 20 | QH | MC | 4 | MCJ |
| 86 | TB | F | 11 | MCJ | 76 | TB | M | 4 | ACJ |
| 105 | Saddlebred | MC | 5 | ACJ | 80 | STB | M | 2 | MCJ |
| Range | | **41% MC, 7% M 51% F** | 1-15 | | 74 | TB | MC | 7 | ACJ |
| Median | | | 5 | | 51 | TB | F | 1 | MCJ |
| Mean | | | 5.24 | | 72 | TB | M | 3 | MCJ |
| SE | | | 0.52 | | 69 | QH | F | 10 | MCJ |
| IQR | | | 2.4 | | 35 | TB | MC | 4 | ACJ |
| | | | | | 39 | TB | MC | 6 | MCJ |
| | | | | | 37 | TB | MC | 3 | ACJ |
| | | | | | 42 | TB | M | 2 | MCJ |
| | | | | | 82 | TB | F | 5 | MCJ |
| | | | | | 43 | TB | F | 2 | ACJ |
| | | | | | 78 | TB | F | 3 | MCJ |
| | | | | | 73 | STB | MC | 5 | MCJ |
| | | | | | 29 | Mixed | MC | 15 | Stifle |
| | | | | | 62 | TB | F | 3 | MCJ |
| | | | | | 44 | QH | MC | 20 | ACJ |
| | | | | | 46 | TB | F | 3 | MCJ |
| | | | | | 54 | TB | F | 2 | ACJ |
| | | | | | 2 | TB | MC | 4 | MCJ |
| | | | | | 60 | TB | F | 4 | ACJ |
| | | | | | 3 | TB | M | 2 | MCJ |
| | | | | | 47 | TB | F | 3 | ACJ |
| | | | | | 49 | QH | MC | 9 | MCJ |
| | | | | | 7 | TB | F | 2 | MCJ |

<table>
<tr><td>84</td><td>TB</td><td>F</td><td>19</td><td>MCJ</td></tr>
<tr><td>28</td><td>TB</td><td>F</td><td>12</td><td>MCPJ</td></tr>
<tr><td>36</td><td>TB</td><td>F</td><td>3</td><td>Carpus</td></tr>
<tr><td>61</td><td>TB</td><td>F</td><td>3</td><td>MCJ</td></tr>
<tr><td>85</td><td>TB</td><td>F</td><td>8</td><td>ACJ</td></tr>
<tr><td>83</td><td>TB</td><td>F</td><td>22</td><td>ACJ</td></tr>
<tr><td>52</td><td>TB</td><td>F</td><td>2</td><td>MCJ</td></tr>
<tr><td>68</td><td>STB</td><td>M</td><td>5</td><td>MCJ</td></tr>
<tr><td>56</td><td>TB</td><td>F</td><td>2</td><td>MCJ</td></tr>
<tr><td>48</td><td>TB</td><td>M</td><td>5</td><td>MCJ</td></tr>
<tr><td>38</td><td>STB</td><td>F</td><td>2</td><td>MCJ</td></tr>
<tr><td>64</td><td>TB</td><td>F</td><td>4</td><td>MCJ</td></tr>
<tr><td>79</td><td>TB</td><td>F</td><td>3</td><td>MCJ</td></tr>
<tr><td colspan="2">Range</td><td rowspan="5">41% MC<br>13% M<br>46% F</td><td colspan="2">0.7-22</td></tr>
<tr><td colspan="2">Median</td><td colspan="2">3</td></tr>
<tr><td colspan="2">Mean</td><td colspan="2">5.09</td></tr>
<tr><td colspan="2">SE</td><td colspan="2">0.52</td></tr>
<tr><td colspan="2">IQR</td><td colspan="2">3</td></tr>
</table>

Table S3. Demographic details of the canine cohort.

Canine synovial fluid samples were obtained from dogs with PTOA of the stifle joint and healthy controls. Values include sample ID, breed, sex, and age. Summary statistics (range, median, mean, SE, and IQR) are provided. Abbreviations: MI, male intact; MN, male neutered; FS, female spayed; FI, female intact.

| **Control** | | | | **OA** | | | |
|---|---|---|---|---|---|---|---|
| No. | Breed | Sex | Age | No. | Breed | Sex | Age |
| 28 | Beagle | MI | 4 | 45 | Mixed Dog | FS | 5 |
| 30 | Beagle | MI | 2 | 55 | Mixed Dog | FS | 6 |
| 67 | Beagle | MI | 3.3 | 52 | Labrador Retriever | MN | 4.2 |
| 27 | Beagle | MI | 4 | 19 | American Pit Bull Terrier | FS | 2 |
| 68 | Beagle | MI | 6.7 | 4 | Labrador Retriever | FS | 8 |
| 71 | Beagle | MI | 7.5 | 38 | German Shepherd | MN | 6.1 |
| 69 | Beagle | MI | 6.8 | 39 | Mixed Dog | FS | 10.1 |
| 70 | Beagle | MI | 4.8 | 63 | Staffordshire Bull Terrier | FS | 7.4 |
| 29 | Beagle | MI | 4 | 48 | Pembroke Welsh Corgi | MN | 3.6 |
| Range | | | 2-7.5 | 33 | Labrador Retriever | MN | 3.3 |
| Median | | | 4 | 46 | Bulldog, English | FI | 5 |
| Mean | | **100% MI** | 4.8 | 65 | Mixed Dog | MN | 9.7 |
| SE | | | 0.6 | 17 | Mixed Dog | MN | 2 |
| IQR | | | 1.45 | 13 | Border Collie | FS | 7 |
| | | | | 40 | Rottweiler | FS | |
| | | | | 61 | English Springer Spaniel | MN | 9.6 |
| | | | | 47 | Labrador Retriever | MN | 4.2 |
| | | | | 11 | Rottweiler | MN | 2 |
| | | | | 64 | Mixed Dog | FS | 11 |
| | | | | 9 | Cane Corso | MN | 1 |
| | | | | 37 | Great Dane | MN | 5 |
| | | | | 56 | Labrador Retriever | | 10.6 |
| | | | | 36 | Rottweiler | MN | 5.3 |
| | | | | 18 | Mixed Dog | FS | 4 |
| | | | | 7 | Mixed Dog | MN | 8 |
| | | | | 15 | Labrador Retriever | MN | 6 |

| | | | |
|---|---|---|---|
| 5R | Mixed Dog | MN | 8 |
| 44 | Labrador Retriever | MN | 4.4 |
| 3 | Mixed Dog | MN | 7 |
| 2R | Rottweiler | FS | 2 |
| 16 | Rottweiler | FS | 2 |
| 53 | Cane Corso | MN | 2.6 |
| 12 | Labrador Retriever | MN | 2 |
| 5 | Mixed Dog | MN | 8 |
| 58 | Labrador Retriever | FS | 10 |
| 49 | Golden Retriever | FS | 4.3 |
| 62 | Mixed Dog | MN | 8.2 |
| 2 | Rottweiler | FS | 2 |
| 6 | Belgian Malinois | FS | 6 |
| 66 | Labrador Retriever | FS | 7.1 |
| 8 | Golden Retriever | FS | 7 |
| 20 | Mixed Dog | FS | 13 |
| 1 | Mixed Dog | FS | 8 |
| 14 | Mixed Dog | FS | 3 |
| 42 | Labrador Retriever | MN | 4.1 |
| 10 | English Setter | MN | 9 |
| 60 | Bernese Mountain Dog | MN | 6.3 |
| 51 | Labrador Retriever | FS | 5.4 |
| 50 | Mixed Dog | MN | 3.5 |
| 59 | Golden Retriever | FS | 9.3 |
| 57 | Golden Retriever | FS | 6.9 |
| Range | | **50% MN 0% MI 48% FS 2% FI** | 1-13 |
| Median | | | 6 |
| Mean | | | 5.90 |
| SE | | | 0.41 |
| IQR | | | 4.4 |

Table S4: Lectins and antibodies highlighted in volcano plots

Table lists the lectins/antibodies annotated on the volcano plots, showing their abbreviated labels (as used in the figures), full names, species of origin, source/vendor, and reference.

| Figure Label | Full name | Origin | Source | Reference |
|---|---|---|---|---|
| LcH$^1$ | Lens culinaris agglutinin | Lens culinaris | Medicago | Bojar et al., 2022 |
| LcH$^2$ | Lens culinaris agglutinin | Lens culinaris | EY | Bojar et al., 2022 |
| anti-LeA$^1$ | Anti-Lewis A | mouse | Abcam | https://www.abcam.com/en-us/products/primary-antibodies/blood-group-lewis-a-antibody-spm-522-ab231432?srsltid=AfmBOorjMm0ilNCjmjLNKNVv9GTOnOTtDsMwPyg6ISHCJcXo4yViRFHR |
| anti-LeA$^2$ | Anti-Lewis A | mouse | Sigma | https://www.sigmaaldrich.com/US/en/product/sigma/sab4700762#product-documentation |

Table S5. Detailed regression results for lectins associated with osteoarthritis (OA) in canine, equine, and human synovial fluid samples with p ≤ 0.05.

Regression coefficients (β) and p values are reported for each lectin. Epitope binding specificities are provided for reference.

Abbreviations: OA, osteoarthritis; β, regression coefficient.

| Species | β coefficient | p value | Lectin | Epitope |
|---|---|---|---|---|
| Dog | -9.90E-01 | 1.54E-02 | ama_vector | Oligo mannose |
| Dog | -4.38E-01 | 4.58E-02 | aol_tci_america | Fucose |
| Dog | 6.40E-01 | 3.96E-02 | anti_h2_scbt | Blood group H2 antigen |
| Dog | -8.81E-01 | 5.32E-03 | anti_lewis_a_abcam | Lewis B |
| Dog | -1.00E+00 | 1.13E-03 | anti_lewis_a_sigma | Lewis B |
| Dog | -8.89E-01 | 4.43E-04 | anti_sialyl_lewis_x | Sialyl Lewis A |
| Dog | 6.98E-01 | 1.92E-02 | bpl_vector | β-Gal / β-GalNAc |
| Dog | 7.78E-01 | 1.16E-03 | ban_lec_h84t_u_alberta | Mannose |
| Dog | -1.03E+00 | 1.65036974151649e-05 | dsa_vector | LacNAc |
| Dog | 5.86E-01 | 6.72E-03 | griffithsin_u_alberta | Mannose |
| Dog | -8.00E-01 | 4.34E-04 | hpa_sigma | Blood Group A |
| Dog | -1.03E+00 | 5.36762610749245e-06 | lc_h_aniara | Core Fucose |
| Dog | -3.85E-01 | 1.36E-02 | pha_e_vector | Bisecting GlcNAc |
| Dog | -9.16E-01 | 9.81315750626087e-06 | psa_glyco_matrix | Core Fucose |
| Dog | -8.43E-01 | 1.51E-02 | psa_vector | Core Fucose |
| Dog | -9.13E-01 | 8.90684613079386e-05 | psl1a | α2,6 sialylation |
| Dog | 7.49E-01 | 2.45E-02 | rca120_vector | Gal / Lac |
| Dog | -1.25E+00 | 2.47851975424988e-07 | slbr_n_u_alberta | α2,3 sialylation |
| Dog | -5.42E-01 | 4.87E-02 | sna_i_ey | α2,6 sialylation |
| Dog | 8.76E-01 | 5.59E-04 | sna_ii_ey | α2 Fucose /oligo mannose |
| Dog | -1.18E+00 | 4.90E-04 | sna_vector | α2,6 sialylation |
| Dog | 8.48E-01 | 2.50E-02 | tja_ii_aniara | α2 Fucose |

| | | | | |
|---|---|---|---|---|
| Dog | 9.60E-01 | 1.81503922213713e-05 | tja_ii_medicago | α2 Fucose |
| Dog | -7.49E-01 | 5.96603119373764e-05 | wga_vector | GlcNAc |
| Horse | -3.75E-01 | 1.69E-02 | aal_medicago | Fucose |
| Horse | 5.30E-01 | 1.45E-02 | aia_ey | β1,3-GalNAc |
| Horse | 7.22E-01 | 9.30708748386415e-05 | aia_glyco_matrix | β1,3-GalNAc |
| Horse | 6.87E-01 | 8.35E-04 | aia_vector | β1,3-GalNAc |
| Horse | -3.71E-01 | 2.41E-02 | anti_lewis_a_sigma | Lewis B |
| Horse | -3.84E-01 | 2.84E-04 | anti_sialyl_lewis_x | Sialyl Lewis A |
| Horse | 4.35E-01 | 3.38E-04 | ban_lec_h84t_u_alberta | Mannose |
| Horse | -3.17E-01 | 4.72E-03 | ca_ey | Bi-antennary N-linked glycans |
| Horse | 3.54E-01 | 9.00E-04 | griffithsin_u_alberta | Mannose |
| Horse | -6.35E-01 | 2.67483605580521e-05 | ltl_lotus_ey | Fucose |
| Horse | -2.41E-01 | 7.97E-03 | lc_h_aniara | Core Fucose |
| Horse | -4.82E-01 | 4.86E-04 | maa_i | Sialylation/Sulfation |
| Horse | -4.41E-01 | 3.19E-03 | maa_i_vector | Sialylation/Sulfation |
| Horse | -3.32E-01 | 3.10E-03 | maa_i_ey | Sialylation/Sulfation |
| Horse | -5.36E-01 | 1.10227309774983e-05 | mal_i_vector | Sialylation/Sulfation |
| Horse | 4.92E-01 | 2.37E-03 | mal_ii | Sialylation/Sulfation |
| Horse | 6.78E-01 | 7.14383802760622e-05 | mpl_vector | β1,3-GalNAc |
| Horse | 2.99E-01 | 8.44E-03 | rca120_vector | Gal / Lac |
| Horse | 3.65E-01 | 4.99E-02 | slbr_b_u_alberta | α2,3 sialylation |
| Horse | 6.84E-01 | 3.62697431588948e-06 | slbr_h_u_alberta | α2,3 sialylation |
| Horse | -3.15E-01 | 7.17E-03 | sna_i_ey | α2,6 sialylation |
| Horse | 4.46E-01 | 4.29E-02 | tja_ii_aniara | α2 Fucose |
| Horse | 3.59E-01 | 1.23E-02 | uda_ey | GlcNAc / Oligo mannose |
| Horse | -3.78E-01 | 3.33E-02 | vva_vector | Terminal GalNAc |

| | | | | |
|---|---|---|---|---|
| Horse | -8.33E-01 | 3.08606967789355 e-05 | wfa_vector | GalNAc-β1,4 |
| Horse | -2.74E-01 | 9.91E-03 | wga_vector | GlcNAc |
| Horse | 6.67E-01 | 2.15E-04 | di_cbm40_u_alberta | _2,3 sialylation |
| Human | -7.27E-01 | 1.72E-02 | aia_vector | β1,3-GalNAc |
| Human | -7.36E-01 | 2.43E-02 | bpl_vector | β-Gal / β-GalNAc |
| Human | 6.99E-01 | 7.92E-03 | con_a_vector | Tri-mannose core |
| Human | 7.44E-01 | 4.26E-03 | griffithsin_u_alberta | High Mannose |
| Human | -8.78E-01 | 2.75E-02 | hpa_sigma | Blood Group A |
| Human | 3.70E-01 | 2.78E-02 | lc_h_aniara | Core Fucose |
| Human | -7.51E-01 | 8.52E-03 | mna_g_ey | GalNAc |
| Human | -7.41E-01 | 1.69E-02 | mpl_vector | β1,3-GalNAc |
| Human | 5.13E-01 | 1.02E-02 | pha_e_vector | Bisecting GlcNAc |
| Human | 3.76E-01 | 3.97E-02 | slbr_n_u_alberta | α2,3 sialylation |
| Human | 6.46E-01 | 1.83E-02 | tl_1_ey | GlcNAc |
| Human | -8.67E-01 | 1.59E-02 | vva_vector | Terminal GalNAc |

Table S5. Performance of machine learning models trained with PyCaret on synovial fluid lectin data.

Model performance metrics are reported as mean values after stratified resampling with Synthetic Minority Oversampling Technique (SMOTE). Metrics include accuracy, area under the receiver operating characteristic curve (AUC), recall, precision, F1 score, Cohen's kappa, Matthews correlation coefficient (MCC), and training time (TT, in seconds).

Abbreviations: AUC, area under the curve; MCC, Matthews correlation coefficient; SMOTE, Synthetic Minority Oversampling Technique; TT, training time.

| Model | Accuracy | AUC | Recall | Prec. | F1 | Kappa | MCC | TT (Sec) | Dataset |
|---|---|---|---|---|---|---|---|---|---|
| Extra Trees Classifier | 0.9283 | 0.9849 | 0.9155 | 0.95 | 0.9276 | 0.8568 | 0.8655 | 0.013 | SMOTE |
| Light Gradient Boosting Machine | 0.9186 | 0.9668 | 0.8782 | 0.9624 | 0.9129 | 0.8378 | 0.8487 | 0.134 | SMOTE |
| CatBoost Classifier | 0.9086 | 0.9842 | 0.8573 | 0.9652 | 0.9007 | 0.8174 | 0.8323 | 0.595 | SMOTE |
| Extreme Gradient Boosting | 0.9083 | 0.9619 | 0.8764 | 0.9431 | 0.9026 | 0.8169 | 0.8272 | 0.01 | SMOTE |
| Random Forest Classifier | 0.899 | 0.9768 | 0.8582 | 0.9465 | 0.8924 | 0.7986 | 0.8132 | 0.015 | SMOTE |
| Gradient Boosting Classifier | 0.8605 | 0.9527 | 0.81 | 0.9125 | 0.8473 | 0.722 | 0.7398 | 0.022 | SMOTE |
| Logistic Regression | 0.851 | 0.9178 | 0.79 | 0.9176 | 0.8326 | 0.7013 | 0.7233 | 0.18 | SMOTE |
| Quadratic Discriminant Analysis | 0.8507 | 0.9609 | 1 | 0.7769 | 0.8728 | 0.7001 | 0.7363 | 0.004 | SMOTE |
| Ada Boost Classifier | 0.8417 | 0.9297 | 0.7909 | 0.9002 | 0.8289 | 0.6842 | 0.7043 | 0.01 | SMOTE |

| | | | | | | | | | |
|---|---|---|---|---|---|---|---|---|---|
| Ridge Classifier | 0.8317 | 0.8942 | 0.75 | 0.9171 | 0.8026 | 0.6624 | 0.6911 | 0.003 | SMOTE |
| Linear Discriminant Analysis | 0.8169 | 0.8799 | 0.73 | 0.9107 | 0.7858 | 0.6326 | 0.6655 | 0.004 | SMOTE |
| K Neighbors Classifier | 0.8021 | 0.9245 | 0.6473 | 0.9514 | 0.7538 | 0.6066 | 0.648 | 0.101 | SMOTE |
| SVM - Linear Kernel | 0.7974 | 0.8808 | 0.7518 | 0.8462 | 0.7844 | 0.5952 | 0.6132 | 0.004 | SMOTE |
| Decision Tree Classifier | 0.7736 | 0.7755 | 0.7236 | 0.8168 | 0.7606 | 0.5486 | 0.5605 | 0.088 | SMOTE |
| Naive Bayes | 0.7583 | 0.8505 | 0.7518 | 0.7931 | 0.7524 | 0.5174 | 0.5436 | 0.004 | SMOTE |
| Dummy Classifier | 0.4833 | 0.5 | 0.6 | 0.2929 | 0.3935 | 0 | 0 | 0.004 | SMOTE |

Table S6: Lectin epitope data

Lectins used for synovial fluid glycomic profiling are listed with species or origin, print concentration, approximate specificity (inhibitory monosaccharide or glycan motif), and vendor/source.

Abbreviations: conc., concentration; EY, EY Laboratories; TCI, TCI America; Mab, monoclonal antibody.

| **Lectin** | **Species/Origin** | **Print Conc.** (μg/mL) | **Rough Specificity /Inhibitory monosaccharide** | **Vendor/Source** |
|---|---|---|---|---|
| AAL | *Aleuria aurantia* | 1000 | Fucose | Vector |
| ACA | *Amaranthus Caudatus* | 1000 | Gal-β1,3-GalNAc | Vector |
| AIA | *Artocarpus integrifolia* | 500 | β1,3-GalNAc | Vector/EY |
| AMA | *Allium moly* | 500 | Oligo mannose | EY |
| Anti-B.G.H2 | MAb mouse IgM [A46-B/B10] | undiluted | Blood group H2 antigen | Santa Cruz Biotechnology |
| Anti-Forssman | MAb Rat IgM [117C9] | undiluted | Forssman Antigen | Abcam |
| Anti-Lewis B | IgM [T218] | undiluted | Lewis B | Sigma |
| Anti-Lewis X | MAb mouse IgM [P12] | undiluted | Lewis X | Abcam |
| Anti-Lewis Y | MAb mouse IgM [F3] | undiluted | Lewis Y | Abcam |

| | | | | |
|---|---|---|---|---|
| Anti-MUC5AC human | Mab mouse IgG1 [CLH2] | undiluted | human MUC5AC | Sigma |
| Anti-MUC5AC mouse | Goat polyclonal to mouse MUC5AC | undiluted | mouse MUC5AC | LSBio |
| Anti-Mucin 15 | Mab mouse IgG1 [H-5] | undiluted | Mucin 15 | Santa Cruz Biotechnology |
| Anti-Sialyl Lewis A | Mab mouse IgG1 | undiluted | Sialyl Lewis A | Abcam |
| Anti-Sialyl Lewis X | Mab mouse IgM | undiluted | Sialyl Lewis X | Abcam |
| AOL | *Aspergillus oryzae* | 1000 | Fucose | TCI America |
| APA | *Abrus precatorius* | 500 | Gal-β1,3-GalNAc / Lac | EY |
| ASA | *Allium sativum* | 1000 | Mannose | EY |
| Blackbean | *Blackbean crude* | 1000 | GalNAc | EY |
| BPA | *Bauhinia purpurea* | 500 | β-Gal / β-GalNAc | Vector |
| BR6 | unknown (from unpublished work) | 480 | under investigation | Gift from Dr. Barbara Bensing |
| CA | *Colchicum autumnale* | 1200 | Bi-antennary N-linked glycans | EY |
| CAA | *Caragana arborescens* | 1000 | Bi-antennary N-linked glycans | EY |

| | | | | |
|---|---|---|---|---|
| Calsepa | *Calystegia sepium* | 1000 | Bisecting N-linked glycans | EY |
| CCA | *Cancer antennarius* | 1000 | 9-O-Acetly sialylation / 4-O-Acetyl sialylation | EY |
| Cholera Toxin | *Vibrio cholerae* | 1000 | GM1 ganglioside | Sigma |
| Con A | *Canavalia ensiformis* | 1000 | Tri-mannose core | EY/Vector |
| CSA | *Cystisus scoparius* | 1000 | Terminal GalNAc | EY |
| DBA | *Dolichos Biflorus* | 1000 | GalNAc | Vector |
| diCBM40 | engineered NanI from *Clostridium perfringens* | 1000 | α Sialylation | Generated in house |
| DSA | *Datura stramonium* | 500 | LacNAc | EY/Vector |
| ECA | *Erythrina cristagalli* | 1000 | LacNAc | Vector |
| EEL/EEA | *Eunonymus europaeus* | 1000 | Blood Group B | Vector/EY |
| GafD | recombinant GafD from *Escherichia coli* | 1000 | GlcNAc | Generated in house |
| GHA | *Glechoma hederacea* | 500 | GalNac | EY |
| GNA/GNL | *Galanthus nivalis* | 1500 | Oligo mannose | Vector/EY |

| | | | | |
|---|---|---|---|---|
| GS-I | *Griffonia simplicifoia-I* | 1000 | α-Gal / Lac | Vector/EY |
| GS-II | *Griffonia simplicifoia-II* | 1000 | GlcNAc | Vector |
| GS-IB4 | *Griffonia simplicifoia-I, isolectin B4* | 2000 | Gal | Vector |
| H84T | *Banana lectin* | 1000 | High mannose | Gift from Dr. David Markovitz |
| HAA | *Homarus americanus* | 1000 | Terminal GalNAc | EY |
| HHL | *Hippeastrum Hybrid* | 1500 | Oligo/High mannose | Vector |
| HPA | *Helix pomatia* | 1000 | Blood Group A | Sigma/EY |
| LAA | *Laburnum alpinum* | 900 | GlcNAc | EY |
| LBA | *Phaseolus lunatus* | 1000 | Blood Group A | EY |
| LcH | *Lens Culinaris* | 1000 | Core Fucose | Vector |
| LEA/LEL | *Lycopersicon esculentum* | 1000 | GlcNAc | Vector/EY |
| LFA | *Limax flavus* | 500 | α Sialylation | EY |
| Lotus | *Lotus tetragonolobus* | 1000 | Fucose | Vector |
| MAA | *Maackia amurensis* | 500 | Sialylation/Sulfation | EY |

| | | | | |
|---|---|---|---|---|
| MAL-I | *Maackia amurensis-I* | 2000 | Sialylation/Sulfation | Vector |
| MAL-II | *Maackia amurensis-II* | 2000 | Sialylation/Sulfation | Vector |
| MNA-G | *Morus nigra Morniga G* | 1000 | GalNAc | EY |
| MNA-M | *Morus nigra Morniga M* | 1000 | Oligo mannose / Gal | EY |
| MPA/MPL | *Maclura pomifera* | 1000 | β1,3-GalNAc | Vector |
| NPA | *Narcissus pseudonarcissus* | 1000 | Oligo mannose | Vector |
| PA-I | *Pseudomonas aeruginosa* | 1000 | Gal | Sigma |
| PA-IL | *bacteria* | 1000 | GalNAc | Generated in house |
| PHA-E | *Phaseolus vulgaris Erythroagglutinin* | 1000 | Bisecting GlcNAc | Vector/EY/Sigma |
| PHA-L | *Phaseolus vulgaris Leukoagglutinin* | 1000 | β1,6 Branching N-Link glycans | Vector/EY/Roche |
| PMA | *Polygonatum multiflorum* | 500 | Oligo mannose | EY |
| PNA | *Arachis hyogaea* | 1000 | Gal-β1,3-GalNAc | Vector/EY |

| | | | | |
|---|---|---|---|---|
| PSA | *Pisum sativum* | 1000 | Core Fucose | Vector |
| PSL | *Polyporus squamosus* | 1000 | α2,6 sialylation | EY |
| PTA | *Psophocarpus tetragonolobus* | 500 | Blood Groups | EY |
| PTL-I | *Psophocarpus tetragonolobus-I* | 1500 | Blood Group A | Vector |
| PTL-II | *Psophocarpus tetragonolobus-II* | 1000 | α2 Fucose | Vector |
| RCA120 | *Ricinus Communis Agglutinin I* | 1000 | Gal / Lac | Vector |
| rCVN | *recombinant Cyanovirin* | 1000 | High mannose | Gift from Dr. Barry O'Keefe |
| rGRFT | *recombinant Griffithsin* | 1000 | High mannose | Gift from Dr. Barry O'Keefe |
| Ricin B Chain | *Ricinus communis* | 1000 | Gal | Vector |
| RPA | *Robinia pseudoacacia* | 500 | Complex N-link glycans | EY |
| rSVN | *recombinant Scytovirin* | 1000 | High mannose | Gift from Dr. Barry O'Keefe |
| SBA | *Glycine max* | 1000 | LacdiNAc | Vector |
| SJA | *Sophora japonica* | 1000 | LacdiNAc | Vector |

| | | | | |
|---|---|---|---|---|
| SK1 | *Streptococcus sanguinis SK1* | 1800 | α2,3 sialylation | Gift from Dr. Barbara Bensing |
| SK678 | *Streptococcus sanguinis SK678* | 450 | α2,3 sialylation | Gift from Dr. Barbara Bensing |
| SLBR-B | *Streptococcus gordonii M99* | 1000 | α2,3 sialylation | Gift from Dr. Barbara Bensing |
| SLBR-H | *Streptococcus gordonii DL1* | 2000 | α2,3 sialylation | Gift from Dr. Barbara Bensing |
| SLBR-N | *Streptococcus gordonii UB10712* | 1000 | α2,3 sialylation | Gift from Dr. Barbara Bensing |
| SNA | *Sambucus nigra* | 500/1000 | α2,6 sialylation | Vector/Sigma |
| SNA-II | *Sambucus nigra-II* | 1000 | α2 Fucose /oligo mannose | EY |
| STA/STL | *Solanus tuberosum* | 500 | GlcNAc | Vector |
| TJA-I | *Trichosanthes japonica-I* | 1000 | α2,6 sialylation | TCI |
| TJA-II | *Trichosanthes japonica-II* | 1000 | α2 Fucose | NorthStar Bioproducts/Aniara Diagnostica |
| TL | *Tulipa sp.* | 700 | GlcNAc | EY |
| UDA | *Urtica dioica* | 1000 | GlcNAc / Oligo mannose | EY |

| | | | | |
|---|---|---|---|---|
| UEA-I | *Ulex europaaeus-I* | 1000 | α2 Fucose | Vector |
| UEA-II | *Ulex europaaeus-II* | 2000 | GlcNAc | Vector |
| VFA | *Vicia faba* | 1000 | GlcNAc | EY |
| VVA | *Vicia villosa* | 1000 | Terminal GalNAc | Vector/EY |
| VVA(man) | *Vicia villosa* | 500 | Mannose | Vector/EY |
| X408 | unknown (from unpublished work) | 1000 | under investigation | Gift from Dr. Barbara Bensing |
| WFA | *Wisteria floribunda* | 1000 | GalNAc-β1,4 | Vector |
| WGA | *Triticum vulgare* | 1000 | GlcNAc | Vector/EY |

Table S7. Lectin microarray workflow and assay conditions.

Detailed workflow for lectin microarray analysis, including sample preparation, lectin library composition, immobilization surface, array production, detection, and data processing.

Abbreviations: PBS, phosphate-buffered saline; PBST, PBS with Tween-20; NHS, N-hydroxysuccinimide; PMT, photomultiplier tube; SNR, signal-to-noise ratio.

| | Description |
|---|---|
| **1. Sample: Glycan-containing sample (e.g. glycan, glycoprotein, cell lysate etc.)** | |
| Description of Sample | Glycoproteins were extracted from hyaluronidase treated synovial fluid from OA affected equine canine and human samples and their healthy controls |
| Sample preparation protocol | Synovial fluid (SF) samples were digested with hyaluronidase at 37°C for 1 hour. Following digestion, 25 µg of protein from each sample was transferred to a 1.5 mL microcentrifuge tube and labeled with NHS-activated Alexa Fluor 555 according to the manufacturer's instructions. A pooled reference sample was prepared by combining aliquots of all individual SF samples and labeling with NHS-activated Alexa Fluor 647. For hybridization, 5 µg of each individual sample and 5 µg of the pooled reference sample were applied to each lectin microarray slide containing more than 100 lectins.<br>Slides were incubated for 2 hours at room temperature in a humidified chamber to allow binding. Following incubation, slides were rinsed gently in wash buffer to remove unbound material, then centrifuged at 500 · g for 5 mins to remove residual liquid. Slides were dried completely and scanned using a Genepix 4400 scanner.<br><br>Data extraction was performed using Genepix Pro 7 software. Probes with a signal-to-noise ratio (SNR) less than 5 in more than 90% of samples were excluded. For all remaining probes, the fluorescence intensity in each channel was normalized to the median of all probe intensities on the array. |

| | |
|---|---|
| Labeling protocol for sample detection | Samples are labelled with Alexa Fluor 555-NHS (Thermo Fisher). |
| Two-color reference (if used) | A pooled reference samples are labelled with Alexa Fluor 647-NHS (Thermo Fisher). |
| Assay protocol | Lectin microarrays are blocked with blocking buffer for one hour at room temperature. Slides are rinsed twice with PBST (0.005%) and once with PBS, then dry the slide using a slide spinner. Each slide was mounted on a 24-well format hybridization cassette (Arrayit), in which each well contains a subarray. To each well, add equal amounts of samples and universal reference, and dilute with PBS and PBST (0.2%) to reach the final volume (150uL). Incubate the slides on an orbital shaker for two hours at room temperature in the dark. After hybridization, wash the arrays with PBST (0.005%) twice for ten minutes, and twice for five minutes. Once finished, remove the slides from the cassette, and immerse the slides in ultrapure water, and dry the slides using a slide spinner. |
| **2. Lectin Library** | |
| General description of the lectin library used in the array | Lectin microarrays are generated in house. |

| | |
|---|---|
| List of lectins and glycan binding proteins, source, concentration and buffer | Please see Table S4. |
| Modification of lectins (e.g. biotin) if any. | N/A |
| 1. **3. Immobilization Surface; e.g., Microarray Slide** | |
| Immobilization surface | Nexterion Slide H Barcoded 3D Hydrogel Coated |
| Manufacturer | Schott North America |
| Custom preparation of surface | N/A |
| **4. Array Production** | |
| Description of Arrayer | Nano-Plotter 2.1 piezoelectric printer (GeSim, Germany) with cooled microwell plate holder and cooled printing deck |
| Lectin deposition | Three replicates of each lectin are printed onto each subarray. |

| | |
|---|---|
| Printing conditions | Dilute lectins to the pre-determined concentrations in the print buffer (final concentration of print buffer: 0.01% Tween-20, 1mM monosaccharide in PBS; Please see **Table S4** for the concentrations of lectins). Load the mixed solution to the microplate. Before printing, check the humidity of the print chamber. The humidity should be kept around 50% during the entire printing. Ensure both microwell plate holder and printing deck are cooled. Adjust the cooling temperature based on ambient temperature and the temperature of the cooled slide deck surface, preventing moisture building up inside the print chamber. Once printing is complete, allow the slides to dry for at least one hour. |
| Array layout | For each microarray, it contains 24 subarrays (3 columns and 8 rows). In each subarray, triplicates of a lectin are printed, and for a row with five lectins, the spot layout should be 15 columns. The row number depends on how many lectin probes are printed on the arrays (i.e., 110 lectins require 22 rows). |
| Quality control | Well-characterized glycoproteins including fetuin, asialofetuin, RNase B and bovine mucin are used for quality control of the printed microarrays. |
| 2. **5. Detector and Data Processing** | |
| Instrument (scanner, flow cytometer) | Fluorescent Slide Scanner Genepix 4300A (Molecular Devices) |

| | |
|---|---|
| Instrument settings | Preview the slide to adjust photomultiplier gain (PMT) for each channel (Alexa Fluor-555: 532nm, Alexa Fluor-647: 635nm) so that the signals are not saturated and within the linear detection range. |
| Image analysis software | GenePix Pro 7 (Molecular Devices) |
| Data processing and statistical analysis | Extracted data is processed for quality checks using Grubbs outlier test with |
| | $\alpha$ = 0.05. Log2 values of the average signals are median-normalized over the individual subarray in each channel. |
| **6. Lectin Microarray Data Presentation** | |
| Data presentation and interpretation | Hierarchical clustering of the processed data is performed using Pearson Correlation coefficient, and visualized with Multi-experiment Viewer (MeV, v4.8, TM4 Microarray Software Suite). If a lectin's SNR (signal-to-noise ratio) < 3 for more than one third of the total samples, then this lectin is considered as inactive and excluded from the list. |

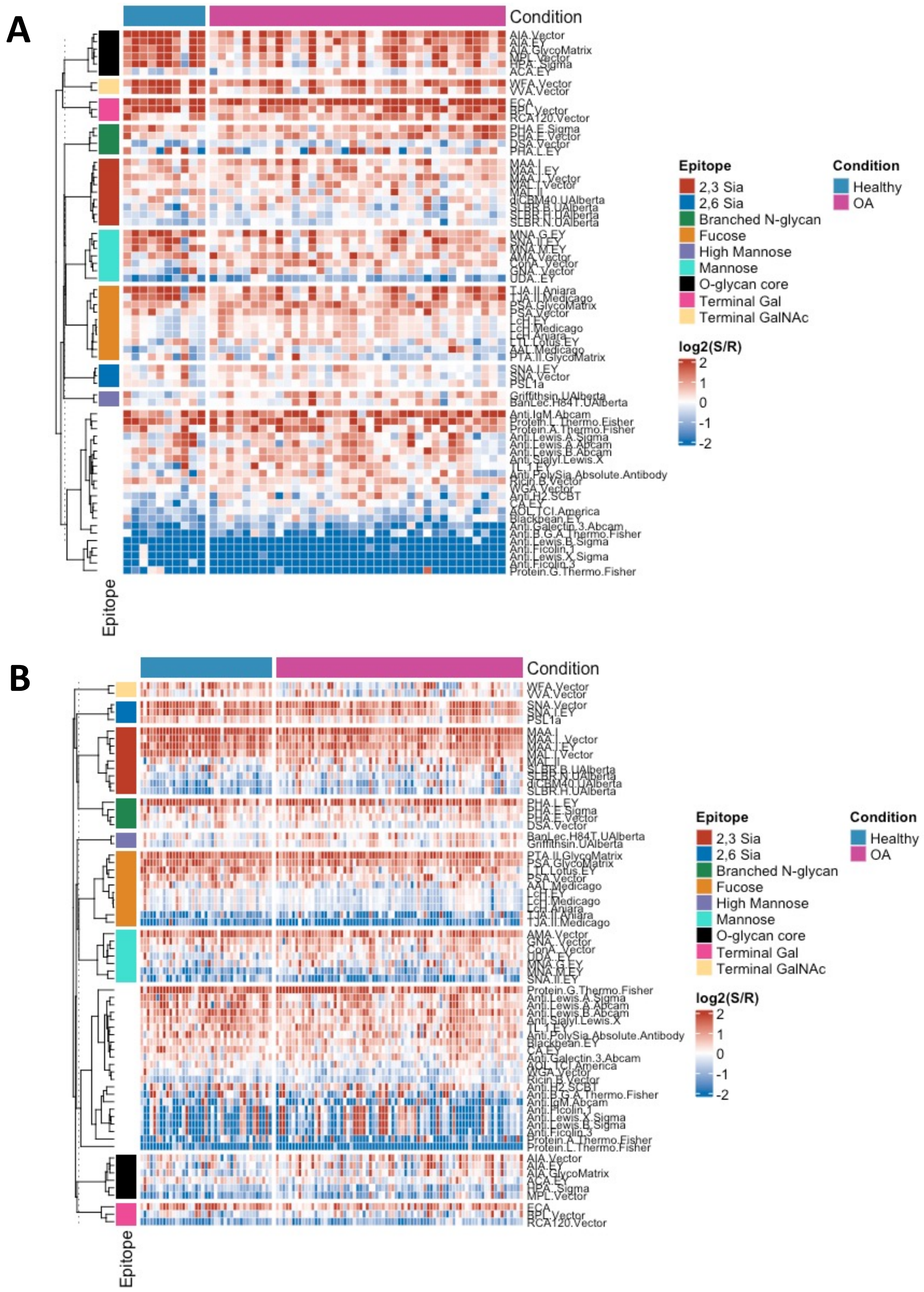
A
B
Condition
Epitope
2,3 Sia
2,6 Sia
Branched N-glycan
Fucose
High Mannose
Mannose
O-glycan core
Terminal Gal
Terminal GalNAc
Healthy
OA
log2(S/R)
2
1
0
-1
-2

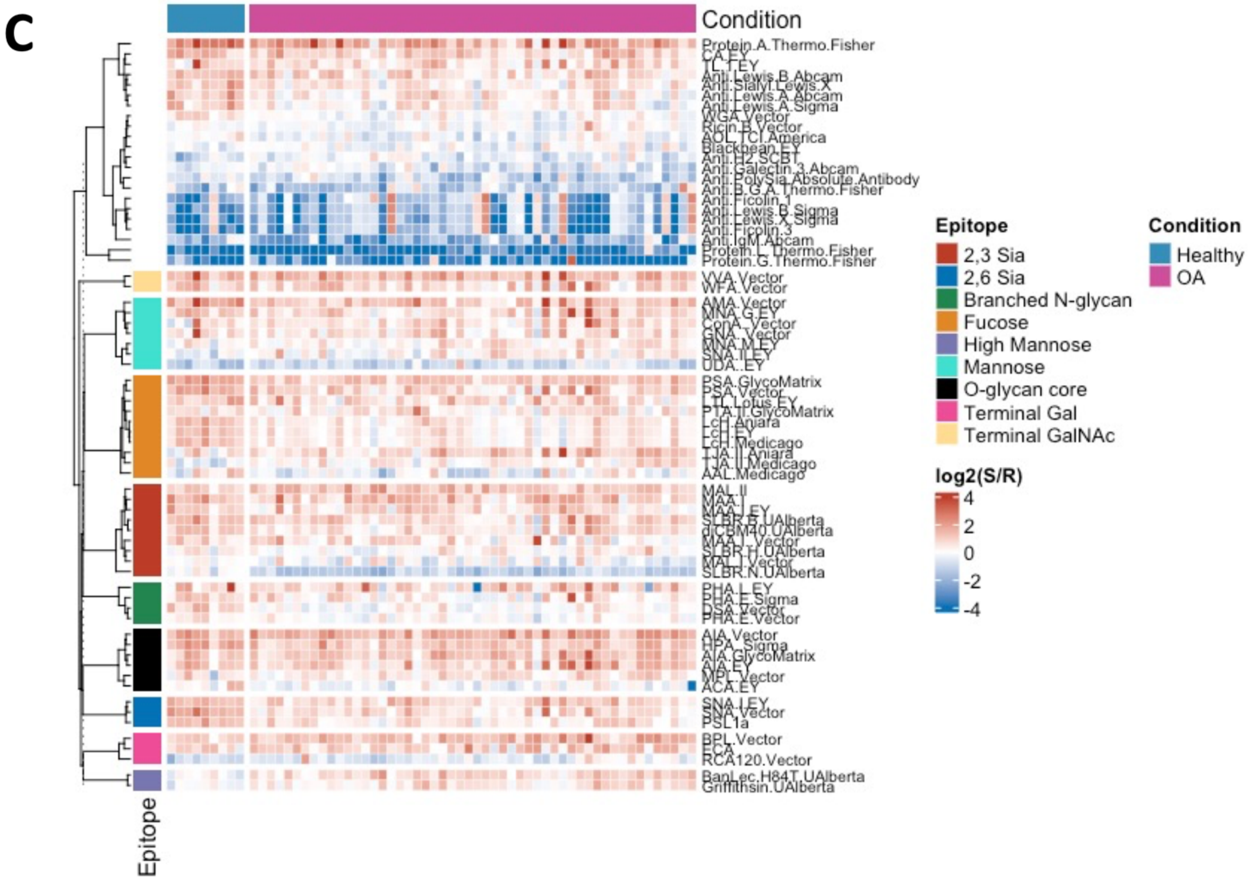


Figure S1. Heatmap of lectin microarray glycopatterns in synovial fluid from human, equine, and canine joints.

(A) Human (knee), (B) equine (carpal), and (C) canine (stifle) synovial fluid samples. Each column represents an individual sample, and each row represents a lectin probe with known glycan specificity (Bojar et al., 2022). Signal intensities are expressed as $log_2$(sample/reference) ratios, with red indicating enrichment and blue indicating depletion relative to the pooled reference. Lectins are grouped by glycan epitope specificity (left color bar). Samples are grouped by condition (top bar: blue = healthy, magenta = OA). Hierarchical clustering was performed using Euclidean distance and complete linkage.

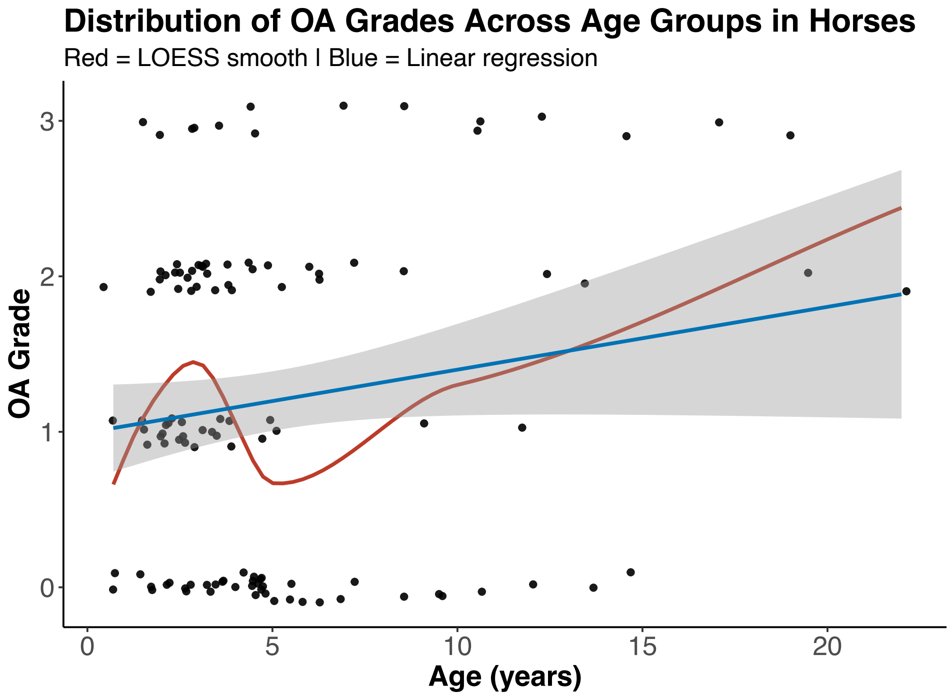


Figure S2. Relationship between age and osteoarthritis grade in equine samples. Scatterplot of age versus OA grade in individual horses. Linear regression (blue) and LOESS fit (red) are shown. Pearson correlation indicated a weak positive trend (r = 0.16, 95% CI: –0.02 to 0.33, *p* = 0.085), while Spearman correlation showed no significant monotonic association (ρ = 0.009, *p* = 0.92).

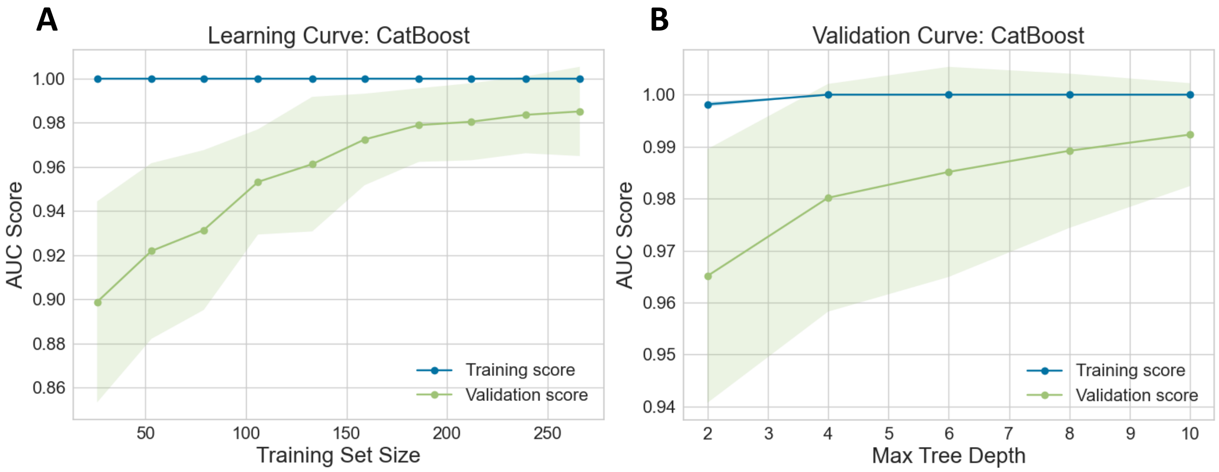


Figure S3. Learning and validation curves for the CatBoost OA classifier.

(A) Learning curve showing training and cross-validation AUC scores as a function of training set size. The classifier shows minimal overfitting and improved generalization with increased training data. (B) Validation curve showing the effect of tree depth on model performance. Deeper trees slightly enhance training AUC, while optimal generalization is achieved at intermediate depths, beyond which validation performance plateaus or declines. Shaded areas represent ±1 standard deviation across cross-validation folds.
